# Functional evaluation of microdissected tumor ‘cuboids’: a microscale *ex vivo* cancer model that retains a complex tumor microenvironment

**DOI:** 10.1101/2024.03.22.586189

**Authors:** Lisa F Horowitz, Ricard Rodriguez-Mias, Marina Chan, Songli Zhu, Yin Tang, Alexandra Locke, Noah R Gottshall, Ivan Stepanov, Casey Stiles, Marcus Yeung, Tran NH Nguyen, Ethan J Lockhart, Charlotte Ahls, Elizabeth Wayne, Wei Wei, Judit Villen, Raymond S Yeung, Taranjit S Gujral, Albert Folch

**Affiliations:** Department of Bioengineering, University of Washington, Seattle, Washington, United States of America; Department of Genome Sciences, University of Washington, Seattle, Washington, United States of America; Human Biology Division, Fred Hutch Cancer Center, Seattle, Washington, United States of America; Institute for Systems Biology, Seattle, Washington, United States of America; Department of Mechanical Engineering, University of Washington, Seattle, Washington, United States of America; Department of Chemical Engineering, University of Washington, Seattle, Washington, United States of America; Department of Biochemistry, University of Washington, Seattle, Washington, United States of America; Department of Surgery, University of Washington, Seattle, Washington, United States of America

## Abstract

To bridge the gap between bench and bedside, there is a need for more faithful models of human cancers that can recapitulate key features of the human tumor microenvironment (TME) and simultaneously facilitate large-scale drug tests. Our recently developed microdissection method optimizes the yield of large numbers of cuboidal microtissues (“cuboids”, ∼(400 µm)^3^) from a tumor biopsy. Here we demonstrate that cuboids from syngeneic mouse tumor models and human tumor biopsies retain a complex TME, making them amenable for drug and immunotherapy evaluation. We characterize relevant TME parameters, such as cellular architecture, cytokine secretion, proteomics profiles, and response to drug panels in multi-well arrays. Despite the cutting procedure and the time spent in culture (up to 7 days), the cuboids display strong cytokine expression and drug responses, including to immunotherapy. Overall, our results suggest that cuboids could provide essential therapeutic information for personalized oncology applications and could help the development of TME-dependent therapeutics and cancer disease models, including for clinical trials.

## Introduction

A fundamental challenge in oncology arises from the small size of most tumor biopsies, in particular for functional drug testing of live tissues *ex vivo*. The amount of tumor material available from any one patient limits both the research and the testing that can be performed from their biopsy. For patients, tumor size at diagnosis is decreasing due to improvements in early detection techniques.^1^ The mean size of tumor resections varies from <1 cm for prostate or breast tumors to up to 9 cm for liver or ovary tumors.^2^ Donated tissue left over from that needed for the patient’s care is typically shared by several investigators, so it is rare for any given lab to obtain more than 1 cm^3^ from a tumor. For mice, maximum volumes are up to ∼0.5-2 cm^3^ for subcutaneous tumors, and often less for internal (orthotopic) tumors due to the greater morbidity. Hence, whether working with mouse or patient tumor, the small amount of starting material limits the scale of drug testing and other studies.

Miniaturization is an obvious solution to the challenge posed by tumor size. Microscale cancer models (µCMs, broadly encompassing organoids, spheroids, tumors-on-chips, tumor slices, fragments, and microdissected tumors) have several advantages over their macroscale counterparts. i) µCMs help maintain *ex vivo* viability by facilitating diffusive mass transport to the inside of the tissue after the resection has interrupted vascular (convective) mass transport.^3^ ii) µCMs can address microscale features of the heterogeneous tumor micro-environment (TME) to properly target and understand the contributions of these unique zones to therapeutic response.^4,5^ iii) µCMs allow for miniaturization of the drug testing workflow.^6^ iv) µCMs enable inexpensive, more efficient tests of high clinical biomimicry.^7^ v) µCMs maximize the use of scarce biopsies.^8–11^ vi) Finally, µCMs minimize animal testing, thus complying with the ethical principles of the 3Rs (Replacement, Reduction, and Refinement).

For patient tumors, functional drug testing addresses the additional challenge from the uniqueness of each solid tumor and the environment in which it grows, determining therapeutic outcomes in ways that cannot be inferred from genetic testing alone.^12^ In the last decade, functional drug testing technologies based on microscale biomimicry have revealed the power and clinical relevance of *ex vivo* testing of cancer patient tissue. These bioengineered non-animal models (NAMs), which help minimize the use of animals, include organs-on-chips^13^ (OOCs) and patient-derived organoids^14^ (PDOs, also termed patient-derived tumor spheroids or organotypic tumor spheroids, among other terms). However, in these NAM platforms, which often take weeks to months to establish, the tumor tissue is generated *de novo* by growing the patient’s cancer cells or by adding separately-derived cells from other patients or from cell lines.^15^ Expansion of the tissue has enabled high-throughput testing with biobanking for many tumors,^14^ and can reveal particular drugs that target key signaling in cancer cells of specific patients.^16^ Tissue bioengineering even allows for building *de novo* microvasculature into the models, e.g., for studying angiogenesis or tumor cell extravasation.^17^ Unfortunately, approaches based on tissue engineering and/or expansion also compromise the TME, including the patient’s immune microenvironment^14,18^ – limiting the relevance of PDOs and OOCs as µCMs, and for immunotherapy in particular since immunotherapy drugs primarily act on the TME.^19,20^ These limitations are a fundamental hurdle for the customization of personalized therapies to the unique tumor and TME of the patient,^21^ and also for the development of immunotherapies (alone and in combinations with other drugs) that often target the human TME.^22^ The pace of progress in clinical studies puts an additional pressure to the development of reliable, quick, and inexpensive drug efficacy tests. Immunotherapy clinical trials, especially in combination with other therapies, are exponentially rising in number.^23^ In sum, traditional bioengineered NAMs like organoids and OOCs allow for high-throughput drug tests but fail to faithfully recapitulate the patient’s TME.

Microdissected tumors (µDTs)^24^ are µCMs based on microdissection, a mechanical approach that separates the tumor into many discrete tissue units without using dissociation enzymes or further expansion. µDTs offer an attractive strategy to preserve the TME intact, maximize the use of scarce biopsies, and minimize the use of animal models.^25,26^ µDTs consist of pieces of tumor that are large enough to preserve the TME and small enough to ensure viability *in vitro* via diffusive transport.^27^ µDTs have been tested since the 1980s in various formats, including precision-cut slices (PCTS)^28–31^ and smaller fragments, which are sometimes referred to as patient-derived explants (PDEs)^26,32–37^ and include patient-derived organotypic spheroids (PDOTS, 40-100 µm-wide).^38,39^ Using scRNAseq, LeBlanc et al. showed that PDEs from glioma could retain much of the spatial heterogeneity of tumors.^40^ PDEs, PCTS, and PDOTs have been used to predict the action of chemotherapeutics^33,35^ and have shown promise for the study of immunotherapy with checkpoint inhibitors,^34,38,41,42^ with some demonstrations of predictability with patient outcomes, but at low throughput.^25,26,31–35^

Various approaches exist for preparation and use of µDTs. Microscale tumor fragments are often created manually by mincing with a scalpel, followed by manual seeding with a pipette and growth in solution or in a gel such as collagen, sometimes on Transwells^43,44^ or in microchannels.^38,39^ Embedding the µDTs in hyaluronan hydrogels enhanced their preservation for up to 10 days.^45^ The use of more precise tools such as a vibratome can standardize the microdissection process. The Gervais group^24,27^ prepared ∼420 µm-diam. cylindrical µDTs by punching cores from patient-derived xenograft tumor slices followed by manual loading into microchannels with traps. Others have performed functional drug tests on microdissected needle core biopsies, e.g., cutting with a vibratome then measuring proliferation changes^46^ or using a proprietary cutting device, culturing in hydrogel and measuring viability and cytokines.^47^ Cordts et al.^48^ developed micromachined silicon blade dicers and stainless steel graters that can quickly generate many tumor fragments. Overall, previous microdissection approaches have successfully demonstrated culture of the tumor in the context of its TME but have often been limited by the poor precision and low throughput of manual procedures.

Motivated by the limited amount of tumor tissue available from each patient, we have recently applied mechanical automation approaches to the µDT workflow. First, we developed a straightforward microdissection technique based on a tissue chopper. With three orthogonal cuts, the chopper mechanically cuts tumors into thousands of regularly-sized, cuboidal-shaped µDTs (referred to as “cuboids”) in ∼30 min.^8^ Due to their small and reproducible size, cuboids can be rapidly arrayed using hydrodynamic traps in microfluidic devices (including a 96-well plate with 4 cuboid traps/well)^8,11^ or loaded by microfluidic ‘lifting’ into multi-well plates using a pipetting robot (up to a 384-well plate, with a programmable number of cuboids/well).^49^ In addition to drug tests by optical readouts,^8,11,49^ we have demonstrated the use of label-free electrochemical aptamers to monitor cell death in multi-well arrays.^50^ However, a rigorous characterization and analysis of the cuboid model was missing.

Here we provide an extensive functional evaluation of mouse and human cuboids as µCMs, including applications not previously explored. We examine the cellular and molecular components of the TME and various readouts in response to drug treatments, including immunotherapy. We also extend the technique to different sizes and formats, including microtissues from core needle biopsies (“discoids”) to improve the clinical applicability of our approach. We characterize the retention and function of the native TME over several days in culture, including cytokine secretion, and demonstrate the addition of exogenous immune cells to contribute to the cuboid TME. In addition to traditional tissue culture platforms, we utilize microfluidic array devices that facilitate 3D imaging and analysis. The regular size of the cuboids enables microfluidic manipulation and analysis, as well as better size selection, e.g., by computer vision and robotics.^49^ We measure traditional viability drug responses and probe deeper into drug responses by targeted protein profiling of signaling pathways and by global quantitative proteomics. Proteomics allows for profiling the TME and for generating a complex readout that encompasses molecular responses to drugs. In sum, cuboids and discoids can serve as a valuable live tumor model with a rich TME. The model supports a wide variety of analytical approaches for deep investigation of drug response for solid tumors in the context of their TME and should complement previous studies of tumor explants, e.g., including those using hydrogel cultures, multiplexed flow cytometry, single-cell sequencing and spatial profiling,^51–54^ that have further supported the potential for intact-TME microtumors to reliably predict clinical responses.

## Results

### Cuboids and discoids as regularly sized microdissected tumor cultures

We microdissected tumors into regularly sized fragments (“cuboids” from surgical biopsies and “discoids” from core needle biopsies) for higher throughput and more precise workflows (see schematic in **Fig 1A**). Robotic, microfluidic, or manual manipulation of the microtumors into wells facilitated culture, drug testing, 3D immunostaining, and analysis. We used multiple mouse and human tumor types to support the generalizability of the approach. For mouse microtumors, we used two cell-derived syngeneic mouse tumor models (with intact immune systems): Py8119 breast cancer (subcutaneous and orthotopic) and MC38 colon cancer (subcutaneous). For patient microtumors, we obtained surgical biopsies from patients with primary liver cancer (intrahepatic cholangiocarcinoma, ICC), from colorectal liver metastases (CRLM), and from metastatic melanoma, all of which had often undergone therapy prior to resection. We chose to focus mainly on 400-µm cuboids for patient and mouse experiments as they have ∼4.1 times more intact tissue volume than the 250-µm cuboids. From previous work by others on tumor spheroids, we inferred that microtumors larger than 400 µm would result in hypoxic cores,^55,56^ so we did not consider larger sizes.

**Fig 1.**
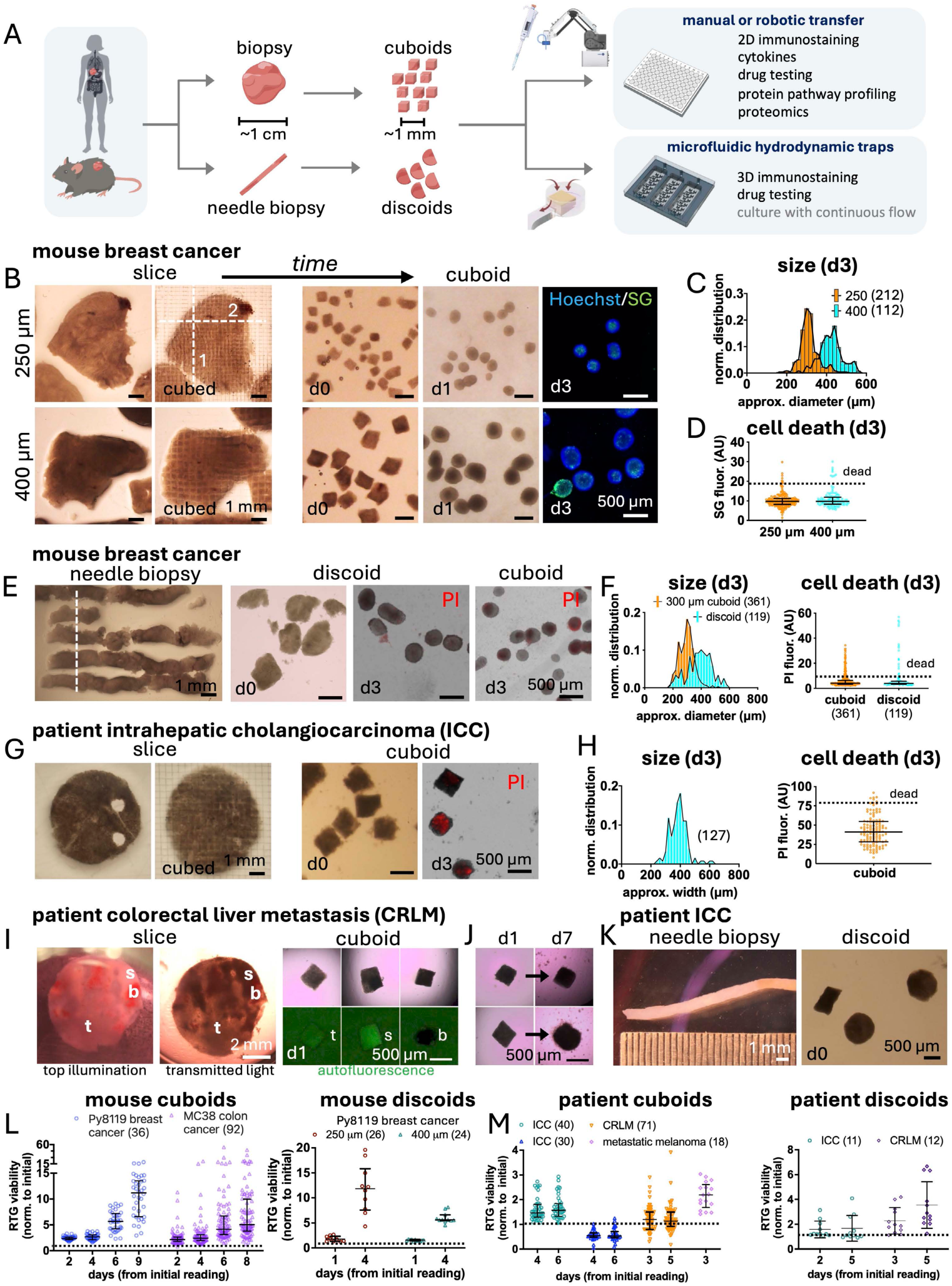
Microtumor generation and culture. (**A**) Workflow for production of microtumors (cuboids from surgical biopsies or discoids from needle cores) and their manipulation (manually, robotically, or microfluidically) for downstream applications. (**B-D**) Formation and culture of 250-µm or 400-µm cuboids from a mouse Py8119 breast cancer syngeneic tumor. Cuboids created by microdissection of vibratome-sectioned slices using a tissue chopper at day 0 (d0), become spheroidal in culture, shown at day 1 (d1) and 3 (d3). Dashed white lines indicate cut orientation and order. Cell death at d3 by SYTOX Green (SG, green, dead nuclei) and Hoechst (blue, all nuclei). (**C**,**D**) Graphs of size and cell death for individual cuboids at d3. Median ± interquartile range. Dashed horizontal line at 2x median. N in parentheses. (**E**) 18-gauge (18G) Py8119 core needle biopsies (left), resulting in 250-µm wide discoids at d0 and d3 (center), and 300-µm cuboids from the same tumor (right). Dashed line indicates tissue chopper cut orientation. Cell death at d3 by propidium iodide (PI, red, dead nuclei). (**F**) Graphs of sizes and cell death at d3 corresponding to (**E**). (**G**) Patient intrahepatic cholangiocarcinoma (ICC) slice and 400-µm cuboids formation and culture (holes secondary to prior needle biopsy extractions). Cell death at d3 (PI). (**H**) Graphs of size and cell death at d3 corresponding to (**G**). (**I,J**) Heterogeneity seen in patient colorectal cancer liver metastasis (CRLM) slices (top illumination or through illumination) and 3 examples of resulting CRLM 400-µm cuboids at d1. t=tumor, s=stroma, b=blood. (**J**) Shape changes seen in 2 representative CRLM cuboids over 6 days. (**K**) Patient ICC 18G needle core biopsy and 400-µm wide discoids. (**L,M**) Sequential RealTime-Glo (RTG) viability graphs of individual microtumors (1 per well) relative to the initial reading. Dashed horizontal lines at viability = 1. (**L**) Mouse cuboids (400 µm) from Py8119 breast tumors and MC38 colon cancer tumors (left). Py8119 mouse discoids (250-µm- or 400-µm wide, right). Initial reading at d1. (**M**) Patient cuboids (left) or discoids (right). Initial reading at d1 except for cuboids from the second ICC and the metastatic melanoma (MM) at d2.

We looked at size and viability in mouse Py8119 breast microtumors for two sizes of cuboids (250 µm and 400 µm) after 3 days in culture (**Fig 1B-D**). We first prepared tumor slices either with a vibratome (250 or 400 µm width) or with a tissue chopper (400 µm) from 6-cm tissue cores, then cut the slices into cuboids by two orthogonal cuts with a tissue chopper (**Fig 1B**), followed by size filtering. Theoretically, an ideal 1 cm^3^ cube of solid tumor would yield 15,625 (25^3^) 400 µm-wide cuboids, or 64,000 (40^3^) 250 µm-wide cuboids assuming three orthogonal cuts and no tissue losses. In practice, depending on the tumor, from an irregular ∼0.5 cm^3^ biopsy we obtained over a thousand ∼400 µm-wide cuboids and thousands of ∼250 µm-wide cuboids, with ∼50% loss of the theoretical yield due to edge effects, tissue variability, and tissue processing (including cutting and size filtering).

Mouse microtumors showed relatively consistent size variability and high viability. We observed that, with cuboids and discoids from mice, the polygonal shape observed immediately after cutting changed to a spheroid shape by the following day. Our size estimates are limited by 2D measurements obtained from images. The estimated diameters of 250-µm cuboids and of 400-µm cuboids were 313 ± 48 µm (AVE ± SD) and 422 ± 58 µm, respectively (**Fig 1B,C**). The diameter of spheroidal-shaped cuboids was estimated from their calculated area on images, assuming a perfect circle. The vibratome, initial tissue conditions, and potential changes during culture (e.g., central necrosis) further contributed to size variability. The tissue chopper itself only contributes a variability of 2-5%, less than a third of the variability seen at day 3 (CV’s between 15-25%). Live/dead staining with the SYTOX Green (SG) nuclear stain for death and the Hoechst blue nuclear stain for all nuclei, showed high death signal in only ∼3% and 7% of 250 and 400-µm cuboids, respectively (**Fig 1D**).

Mouse discoids from needle biopsies shared similar viability with the mouse cuboids. To make discoids (**Fig 1E**), we cut an 18G needle biopsy (838 µm-diam., semi-circular cutting profile) into 250 µm (or 400 µm) wide discoids with one pass of the tissue chopper. While discoids are larger than cuboids, the maximum theoretical yield of only 40 and 25 discoids, respectively, from a 1 cm-long biopsy underscores the need to maximize yield and regularity. In practice, we obtained less than the predicted amount (e.g., ∼50% for 250-µm discoids from a mouse breast cancer needle biopsy), largely due to variations in tissue quality along the biopsy. For 250-µm thick discoids and 300-µm cuboids prepared from the same mouse breast cancer tumor, the average estimated diameters were 377 ± 87 µm and 261 ± 52 µm, respectively (**Fig 1F**, left). Cell death by propidium iodide red nuclear stain (PI) revealed similar percentages (∼17%) of discoids and of 300 µm-cuboids with high death signal (**Fig 1F**, right).

As opposed to mouse cuboids, patient cuboids generally retained their cuboidal shape over days and when they changed shape, such changes were more variable, reduced, and slower (**Fig 1G**). The shape change in the microtumors may result from differences in extracellular matrix (ECM) makeup, as well as differences in cellular properties between the mouse and human tumors. Wu et al. described a similar myosin II-dependent contraction in tissue slices.^52^ For consistency, we refer to both mouse and human microtissues by their initial shape, “cuboids” or “discoids,” irrespective of any subsequent changes in shape.

We also looked at the size and viability of patient microtumors, which we expect may differ from the mouse microtumors due to the heterogeneity in patient tumors and the fast growth of our cell-derived mouse tumors (**Fig 1G-K**). Regional heterogeneity was visible in low magnification brightfield and fluorescent images of patient CRLM slices and cuboids (**Fig 1I,J**) and ICC needle biopsy and discoids (**Fig 1K**), including bloody and stromal/fibrotic regions. This regional heterogeneity likely results from changes in the TME in response to cancer treatments prior to resection.^57^ As we observed in CRLM slices previously,^6^ stromal areas were darker on brightfield and more autofluorescent as compared to tumor-rich regions. Large, clearly bloody, necrotic regions were removed before chopping into cuboids. Note that these patient ICC and CRLM cuboids retained a cuboidal shape (**Fig 1G,I**), with a mean side length for ICC cuboids after 3 days of 380 ± 61 µm (estimated from area). By PI cell death staining, ∼5% of the cuboids had a high death signal. Analysis of cuboid size from another patient tumor (CRLM) showed similar sizes before and after two days in culture (493 ± 62 µm at d0 unselected and 463 ± 47 µm at d2, robot size selection). In sum, despite the higher heterogeneity in the patient tissue, we found that the mouse and human microdissected tumors had similar uniformity in size and low levels of cell death after 3 days in culture.

To confirm the viability of individual microtumors over longer time periods, we used a non-destructive luminescent viability assay, RealTime-Glo (RTG), which is not affected by the autofluorescence often seen in patient tissues (**Fig 1L,M**, **S1 Fig**). We plated defined numbers of microtumors per well in 96 or 384 well plates, generally using custom-programmed robots (see Methods). We periodically measured luminescence with a plate reader starting the day following addition of the RTG reagents, with no medium changes. Graphs of relative changes in viability are shown in **Fig 1L,M** and absolute measurements in **S1 Fig** Individual 400 µm cuboids from Py8119 mouse breast cancer and MC38 mouse colon cancer tumors showed dramatic, 5-10 fold increases in viability signal over a week in culture (**Fig 1L** left), as did two sizes of Py8119 mouse discoids (250 and 400 µm) over 5 days (**Fig 1L** right, **S1 Fig**). The variability in the signal also increased with time, presumably due to differences in growth between cuboids. Py8119 cuboids grown in different media showed steeper growth and more variability in the serum-containing medium (**S1 Fig**). The placement of multiple cuboids per well revealed the time/nutrient limitations of each situation (**S1 Fig**). Experiments in 384-well plates with 1 cuboid/well, Py8119 and MC38 cuboid viability continued to increase, but with 4 cuboids/well signal peaked at 5 days before dropping, presumably due to insufficient nutrients. In contrast, in 96 well plates with 1, 4, and 8 MC38 cuboids per well, viability did not drop, suggesting sufficient medium. With the Py8119 discoids, viability continued to increase over 5 days, with relatively similar readings for both 250- and 400-µm sizes (**S1 Fig**). For cuboids and discoids, an increase in amount of tissue (by size or by number of cuboids) resulted in a relatively small increase in the absolute viability signal. Potential causes include interactions between the tissues, such as growth suppression due to lowered availability of nutrients or the increased levels of secreted signaling molecules. Thus, mouse cuboids and discoids showed robust viability increases when cultured with different formats, with dependence on culture medium and on cuboid number per well.

In contrast to mouse microtumors, RTG viability in patient microtumors was more heterogeneous and did not increase dramatically (**Fig 1M**, **S1 Fig**). This pattern was seen in cuboids (from ICC, CRLM, and metastatic melanoma, MM) (**Fig 1M**, left) and discoids (ICC and CRLM) (**Fig 1M**, right). The relatively higher viability signal increase in mouse microtumors likely reflects the rapid replication rate of the tumor cell lines used to generate the mouse tumors. Graphs of individual patient microtumor absolute viability also revealed greater variability than in mouse, with clear subsets with very low signal without increase (**S1 Fig**). These differences likely reflect the greater heterogeneity of the patient tissue, which can include cell-poor stromal regions. One way to minimize the variability, in particular the lack of signal in some cuboids, is to combine multiple cuboids per well, such as 8 cuboids/well (96-well plate, **S1 Fig**). Together, these RTG and cell death measures confirmed the viability of the cultured microtumors, though they can only provide bulk measurements without regard to cell type.

### The complexity of the mouse cuboid TME

Using multi-immunohistochemistry (multi-IHC), we evaluated spatiotemporal cellular changes within the cuboid TME, which is expected to remodel over time due to cell growth, cell migration, phenotype alterations, and/or cell death.^29^ Sampling over a week in culture represented a sufficient timeframe for functional assays and minimization of changes from culture. We looked at Py8119 mouse breast cancer cuboids of different sizes and different tumor locations: orthotopic (mammary fat pad injection, one tumor with 400-µm cuboids; two tumors with 250 µm and 400-µm cuboids) and subcutaneous (2 tumors with 400-µm cuboids) (**Fig 2A-C**, images of 250 µm in **S2 Fig**). The two locations allowed us to compare cuboids from tumors that shared the same tumor cells but differed in their TME locations. Histology by hematoxylin and eosin (H&E) staining showed general maintenance of tissue structure by H&E and shrinkage by day 7. We quantitated the multi-IHC immunostaining with the image analysis program HALO (**Fig 2B,C**). As we obtained similar results from both tumor locations (plotted individually), we combined their results for mean values. The finding from these studies are detailed below.

**Fig 2.**
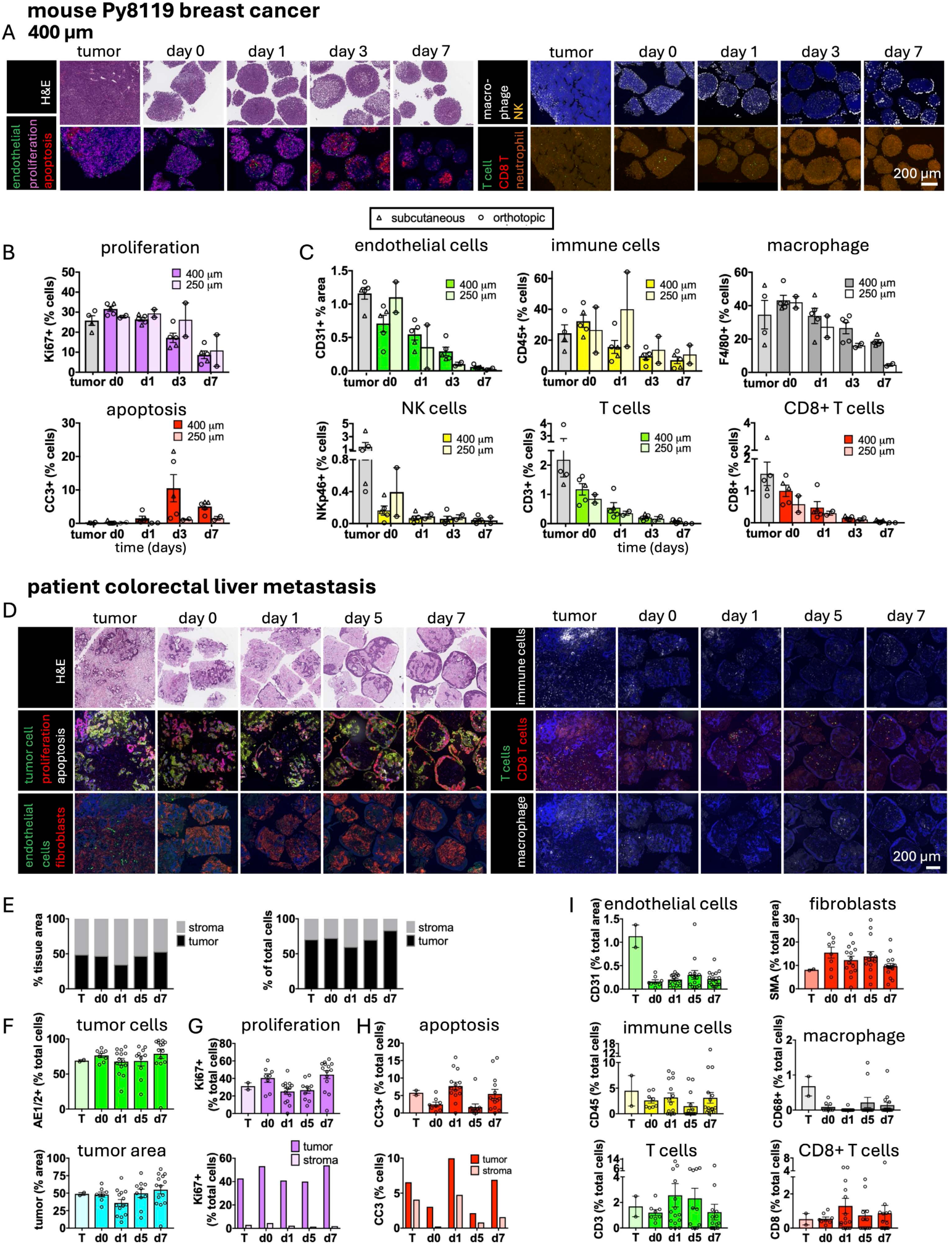
Characterization of cuboids and their TME. (**A-C**) Histology and fluorescent multi-immunohistochemistry (multi-IHC) of cuboids from multiple different Py8119 mouse breast cancer tumors grown in 6-well plates. (**A**) Representative images of histology (H&E) and multi-IHC of 400-µm cuboids from an orthotopic tumor, including apoptotic cell death (cleaved-caspase 3, CC3), proliferation (Ki67), and multiple cell types. (**B,C**) Quantitation of multi-IHC staining of 400-µm cuboids (n=5 and AVE ± SEM) and 250-µm cuboids (n=2, AVE and range) for apoptosis and proliferation (**B**) or cell type (**C**). Individual points represent the mean value for that time point for the total surface area encompassing multiple cuboids (11-40 for 400-µm cuboids) for each tumor. 2 subcutaneous tumors (D) and 3 orthotopic tumors (o). AVE ± SEM. Endothelial cells (CD31), immune cells (CD45), macrophage (F4/80), NK cells (NKp46), T cells (CD3), cytotoxic T cells (CD8), and neutrophils (Ly6G). (**D-I**) Histology and multi-IHC of colorectal cancer liver metastasis (CRLM) 400-µm cuboids cultured in 6-well plates and sampled over a week, as well as the initial tumor (T)). Cell type antibodies: tumor cells (AE1/AE2), endothelial cells (CD31), fibroblasts and smooth muscle cells (smooth muscle actin), immune cells (CD45), macrophage (human: CD68), T cells (CD3), and cytotoxic T cells (CD8). (**E-I**) Quantitation of individual cuboid staining using HALO. Breakdown of staining into “tumor” and “stromal” subregions for overall area in (**E**) or for individual cuboids (**F**), as defined by DAPI and AE1/AE2 staining. (**G**,**H**) proliferation and apoptosis by cuboid and overall distribution in tumor versus stroma. (**I**) Quantitation of cell types by cuboid. AVE ± SEM with individual cuboid values plotted: initial tumor (2), d0 (8), d1 (14), d5 (14), d7 (16). DAPI nuclear stain in blue for mouse and patient immunostaining.

Cuboid health in culture was evidenced by restricted apoptotic cell death and continued proliferation (**Fig 2A,B**, **S2 Fig**). Apoptosis (cleaved-caspase 3, CC3+) in 400-µm cuboids was minimal initially, limited to necrotic regions from the initial tumor seen at day 0 and a few scattered cells at day 1. At day 3, apoptosis peaked (∼10% of the cells), mostly in the central region, then decreased at day 7, whose smaller size likely results from remodeling after cell death. In contrast, the 250-µm cuboids showed minimal apoptosis and did not get smaller over time, as would be expected from improvements in diffusive transport of nutrients and oxygen from the smaller size (**S2 Fig**, **Fig 2B**). Proliferation (Ki-67) in 400-µm cuboids started very high (∼30% at day 0 and day 1) and gradually reduced to ∼9% at day 7. Proliferation trends in 250-µm cuboids were similar. MC38 colon cancer cuboids also showed continued proliferation but differed from Py8119 cuboids by a less dramatic shape change and more areas of necrosis from the initial tumor (present in the initial tumor tissue) (**S2 Fig**). The strong proliferation markers are consistent with the robust RTG viability of Py8119 and MC38 cuboids over a week in culture (**Fig 1**, **S1 Fig**). In sum, immunostaining revealed clear proliferation in both sizes of mouse cuboids, with some central cell death in the first few days for the larger 400-µm cuboids only.

Stromal cell types in the cultured Py8119 cuboids persisted, though with a gradual decline (**Fig 2A,C**; **S2 Fig**). Endothelial cell staining (CD31) in 400-µm cuboids gradually diminished from ∼0.7% of the total area at day 0 to 0.3% at day 3, then to minimal staining at day 7. This rapid disappearance of vascular structures *ex vivo* is a common observation in the absence of circulation,^58^ and we have addressed a potential solution using dynamic-flow cuboid cultures elsewhere.^59^ In contrast, endothelial staining appeared relatively persistent in the MC38 colon cancer cuboids over the week (**S2 Fig**). Staining for immune cells (CD45) was strong initially at ∼32% of total cells, consistent with the previously reported CD45+ fraction in PY8119 tumors (**Fig 2A,C**; **S2 Fig**).^28,60^ The CD45+ percentage gradually diminished, with 17% at day 1, and gradually decreased to 7% at day 7 in 400-µm cuboids.

Particular immune cells are known to play specific roles in tumor biology and response to therapies, such as tumor-associated macrophages (TAM) and cytotoxic T cells.^61^ Macrophages and T cells persisted in the cultured Py8119 cuboids (**Fig 2A,C**; **S2 Fig**). We found that TAMs (F4/80+) were the most frequent immune cells, which is consistent with flow analysis of cultured Py8119 slices^28^ and with a previous study of *in vivo* tumors.^60^ TAMs comprised ∼43% of the total cells in the 400-µm cuboids initially, gradually decreasing to less than half the initial amount at day 7. While there appeared to be abundant tumor-associated neutrophils (Lys6G+), the weak staining made quantitation not feasible. We also observed scattered T cells (CD3+), including some CD8+ T cells, with T cells representing ∼2% of cells at day 1 and gradually diminishing to one tenth of the initial amount by day 7 in 400-µm cuboids. We did not see many natural killer cells (NKp46+) in the cuboids (∼0.07% of cells on average for all days), which is consistent with the few clusters of positive cells seen in sections of the initial tumor. We observed similar trends of immune cells in the 250-µm cuboids through day 7, except for ∼4-fold fewer TAMs (4.5%) at day 7. Multi-IHC of MC38 cuboids over a week in culture qualitatively also showed relatively high levels of macrophage staining and very few T cells throughout (**S2 Fig**). Thus, we detected high levels of macrophages, low levels of T cells, and almost no NK cells in both the initial tumor and in cultured cuboids.

Like any explant, cuboids in culture can be affected by multiple experimental factors. Thus, we also examined cuboids under a few different culture conditions. Culture within collagen affected viability compared to plain medium but did not lead to clear differences by multi-IHC (**S3 Fig**). Cuboids after cryopreservation with two different solutions showed reduced viability by RTG initially, but with strong increases over time, with neither better than the other despite differences in multi-IHC (**S4 Fig**). These data further supported the robustness of the cultured cuboid model with strong viability and a rich TME, and the importance of evaluation of each experimental model for its application.

### The TME and spatial heterogeneity in patient cuboids

In contrast to relatively homogeneous mouse tumor models like above, patient tumors are known to have a lot of spatial heterogeneity (e.g., metabolic, cellular, and genetic), both within tumors and between tumors of the same type.^28^ We observed marked cellular heterogeneity by multi-IHC of 400-µm cuboids from multiple patient tumors: cuboids from a CRLM (**Fig 2D-I**) and a metastatic melanoma from a lymph node (MM) (**S5 Fig**) were studied over a week in culture, and ICC cuboids were studied at day 0 and day 1 (**S5 Fig**). The MM cuboid staining and histology appeared to conform more to the melanoma tumor which had invaded the lymph node than to the normal morphology of a lymph node, though further immunostaining could better distinguish the components and structure of lymph nodes. In general, immunostaining demonstrated the general health of patient cuboids, with low cell death and continued proliferation. Histology and cell type distribution (including fibroblasts, endothelial cells, immune cells, T cells, CD8+ subset of T cells, and macrophages) showed great variability from cuboid to cuboid, as expected from their uneven distribution in the original tumors. In sum, in both tumors followed over a week, cell type staining showed continued presence of all cell types in the cuboids over the week.

For the CRLM cuboids, more in-depth analysis of the staining by individual cuboids allowed us to quantitate the structural and cellular heterogeneity, with the caveats that sampling was limited (though random) and each cuboid was represented by only one cross-section (**Fig 2E-I**). The CRLM tissue and cuboids had subregions of “tumor” (i.e., dominated by tumor cell staining) and stroma (i.e., delineated by fibroblast staining), clearly seen by H&E histology. The CRLM cuboids showed some remodeling over time, but with qualitative maintenance of the tumor microstructure. From day 1 onward the tumor cells appeared to form a superficial layer in addition to the tumor cells inside, consistent with their epithelial origin and their histology in the metastatic tumor. Interestingly, CRLM cuboids grown within collagen for 5 days did not show this remodeling, presumably due to exposure to an ECM-like surface (**S6 Fig**). Collagen also resulted in steadier levels of RTG viability. Remodeling also occurred with cryopreservation of CRLM, with clear cell death by histology (H&E) and RTG viability using several specialized media but followed by growth and recovery (**S7 Fig**). For quantitation, we defined tumor/stroma subregions using tumor cell staining (AE1/2+) and nuclear staining (DAPI) (**Fig 2F-H**). Despite the remodeling, we found that the relative amount of tumor (versus stroma) remained relatively constant over the week, with ∼45% of the area and ∼70% of the cells (**Fig 2E,F**). Therefore, the overall regional tumor-stromal heterogeneity in patient tumors persisted in cuboids, with remodeling in response to liquid culture and not to hydrogel culture.

The CRLM and MM cuboids showed continued proliferation (Ki-67+) and low cell death (CC3+, apoptosis) throughout the week in culture (**Fig 2D,G**,H, **S5 Fig**). In CRLM cuboids, both proliferation and apoptosis localized to tumor regions, with high proliferation (33% of total cells on average, 40% on day 0) and scattered low level baseline apoptosis (5% on average, 3% day 0). However, the central apoptosis we had observed in the mouse cuboids was not apparent in the patient cuboids (CRLM or MM). Indeed, the relatively consistent size of cuboids over the week (in contrast to the decrease in 400-µm mouse cuboids) suggests that central necrosis had not occurred. The more stable cell death and proliferation in human versus mouse cuboids may reflect the lower cell density and cell growth in the human cuboids. In sum, low cell death and continued proliferation in patient cuboids looked similar to the initial tumor and suitable for *in vitro* studies that require proliferation, such as cytotoxic chemotherapy.

In the CRLM cuboids we saw continued presence of endothelial cells and cancer-associated fibroblasts (CAFs), stromal cells known to play important roles in tumor biology and drug responses^62^ (**Fig 2D,I**). Clear endothelial cell staining (CD31+) remained low at ∼0.2% of the total area over the week in contrast to the decrease seen in mouse cuboids (**Fig 2C**). CAFs (smooth muscle actin, SMA) clearly colocalized with the relatively acellular, non-tumor, stromal areas and remained relatively constant from day 0 to 1 week in culture (∼13% of total area) (**Fig 3A,G**). In contrast, in the MM initial tumor and cuboids, SMA staining appeared to mainly demarcate smooth muscle cells that surrounded CD31-positive blood vessels (**S5 Fig**). In conclusion, CAF and endothelial staining in cuboids resembled that in the initial tumors.

**Fig 3.**
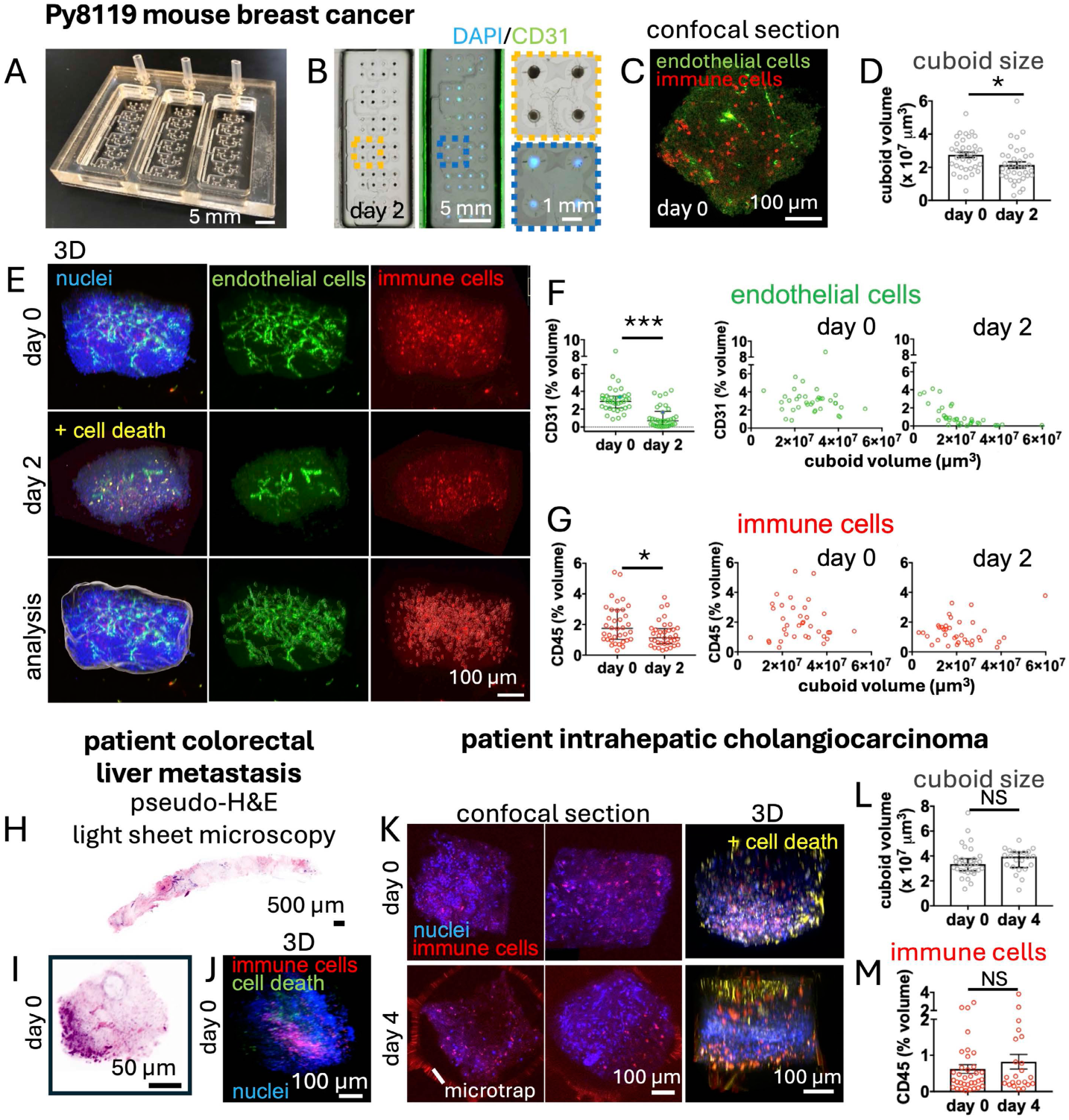
3D analysis of the cuboid TME. (**A-G**) Whole-mount immunostaining of orthotopic Py8119 mouse cuboids at day 0 and day 2. (**A**) Microfluidic device used for on-device immunostaining, clearing, and confocal imaging. Each of 3 independent wells have 48 cuboid traps connected by binaries to a single output. (**B**) One well showing Py8119 mouse cuboids in the device after 2 days in culture and immunostaining (CD31 for endothelial cells, CD45 for immune cells, cleaved-caspase-3 for cell death, DAPI nuclear stain), and clearing with Ce3D. (**C**) Confocal image of immunostaining for day 0. (**D**) Graph of cuboid volume. AVE ± SEM. n=34-36 cuboids. Student’s T-test, *p=0.02. (**E**) 3D projection of cuboids stained in the traps with example of Imaris 3D analysis for day 0. (**F,G**) Graphs showing quantitation of endothelial and immune cell staining versus volume for individual cuboids (points). Median and quartile. n=34-36 cuboids. Mann-Whitney test. *p=0.02, ***p<0.0001. (**H,I**) Optical sections from pseudo-H&E of open top light sheet microscopy of intact needle biopsy (**H**) and cuboid (**I**) from a patient colorectal liver metastasis (CRLM). (**J**) 3D projection of a CRLM cuboid immunostained with CD45 for immune cells, cleaved-caspase-3 for cell death, and a DAPI nuclear stain. (**K**) Patient idiopathic cholangiocarcinoma cuboids immunostained at day 0, or after 4 days culture in the microfluidic device, with representative 2D confocal sections of two cuboids and a 3D projection the second cuboid. (**L,M**) Quantitation of cuboid volume (**L**) and of immune cells (**M**) (% of total cuboid volume). n=34 (day 0) and n=22 (day 4) cuboids. AVE ± SEM. Student’s T-test (volume) and Mann-Whitney test (CD45), NS.

Immune cells were present throughout culture, with clear variability in spatial distribution and over time (**Fig 2D,I**). The non-specific marker for immune cells, CD45, revealed uneven regional localization of immune cells in the initial tumors and in the cuboids (**Fig 2D**), as well as in the MM (**S5 Fig**). Quantitation for immune cells in the CRLM cuboids showed an average of ∼3% of cells total throughout, but with a dramatic decrease in many individual cuboids with culture (**Fig 2I**). In contrast to the Py8119 mouse breast cancer tumor staining, which had a predominance of macrophage with few T cells, the human tumors and cuboids had relatively few macrophages (CD68+) and many T cells (CD3+), with large numbers in some cuboids and few to none in others (**Fig 2D,I**, **S5 Fig**). In both the CRLM and ICC tumor and cuboids, roughly half of the T cells appeared to be CD8+ (**Fig 2I**, **S5 Fig**), while in the melanoma, most of the T cells were CD8+ (**S5 Fig**). Quantitation in the CRLM showed relatively stable levels of T cells (CD3+) with an average of 1-2.5% of total cells overall (**Fig 2I**). The results in the CRLM cuboids were consistent with findings from a 25-plex multi-IHC study of the immune microenvironment in direct analysis of CRLM biopsies.^63^ In sum, over a week in culture, the patient cuboids maintained the tissue complexity and heterogeneity of the initial tumors, including continued cell proliferation and a diversity of cell types.

### 3D analysis of the TME in intact cuboids

Within a tumor, immune and endothelial cells are not distributed homogeneously, but their 3D distribution is difficult to assess from isolated 2D sections. To directly evaluate the 3D distribution and heterogeneity of immune and endothelial cells among individual cuboids, we performed 3D immunostaining, confocal imaging, and analysis of intact 400-µm cuboids from mouse Py8119 breast cancer tumors (**Fig 3A-G**). We performed tissue clearing to improve the limited light penetration into the 400-µm cuboids. For these experiments, we used microfluidic devices with arrays of hydrodynamic cuboid traps (∼600 um diameter, 1 mm depth) that enabled the entire workflow of culture, immunostaining, clearing, and confocal imaging (**S8 Fig**). The device design was a simplified version of a more complex, valved, CNC-milled devices previously used for drug testing.^11^ We fabricated our devices by laser cutting with three different configurations of microtrap arrays. For mouse cuboid immunostaining, the device consisted of three independent rectangular wells each with 48 microtraps connected to one outlet (144 traps total) (**Fig 3A,B**, **S8 Fig**). To load the device, we used negative pressure to pull the cuboids into the traps (see Methods). We confirmed viability for mouse and patient cuboids in the device after loading and culture by IVIS imaging of RTG luminescence (**S8 Fig**).

Confocal imaging of the immunostained/cleared Py8119 mouse cuboids allowed us to simultaneously measure cuboid volume and assess the presence of endothelial cells (CD31+) and immune cells (CD45+) (**Fig 3C-G**, **Movies S1** & **S2**). To consider possible effects of the TME, we studied cuboids from two subcutaneous and two orthotopic Py8119 tumors, however we did not observe any obvious qualitative differences between subcutaneous and orthotopic Py8119 tumors. The cuboids were sampled at day 0 and at day 2, as CD31 staining had mostly disappeared by day 4 (**Fig 3**; Movie **S2**). Qualitative assessment of immunostaining showed abundant CD45+ immune cells at both time points, though less at day 2, and a marked reduction in endothelial cells at day 2 (**Fig 3C,E**). Staining for apoptosis (CC3+) revealed scattered positive cells at day 2 (**Fig 3E**). To better understand the distribution of sizes and immunostaining among individual cuboids, we performed quantitative 3D image analysis of stained structures for one orthotopic tumor using Imaris image analysis (**Fig 3D,F**,G). While the cuboid volume was very variable, the mean volume decreased from day 0 to day 2 (∼23%, **Fig 3D**), consistent with the shape changes seen in 2D images. The endothelial cell staining by day 2 (CD31 volume fraction) was similar to the decrease seen by day 3 using multi-IHC (∼60%, see **Fig 3F** and **Fig 2C**). Comparison with cuboid size revealed that the CD31 decrease occurred in the larger cuboids (**Fig 3F**). The ∼30% decrease in immune cell staining by day 2 (CD45 volume fraction, **Fig 3G**) was consistent with the decreases seen by multi-IHC (**Fig 2C**). However, in contrast to CD31, the CD45 volume fraction did not show a dependence on cuboid size. Thus, after 2 days in culture, CD31 decreased in larger cuboids and not in smaller cuboids, while CD45 decreased in both sizes.

3D imaging of intact CRLM and ICC patient cuboids confirmed the high heterogeneity shown above by 2D immunostaining and viability. We used cuboids from one CRLM tumor (day 0 only) and one ICC tumor (day 0 and day 4). This variability of the histology both at distant regions and locally within the dimension of a single cuboid is clearly seen in whole-mount pseudo-H&E staining and lightsheet microscopy of a CRLM needle biopsy and cuboid (**Fig 3H,I**). In 3D immunostaining experiments on patient cancer cuboids (**Fig 3J,K**; **Movies S3** & **S4**), we looked at immune cell (CD45+) distribution as well as at apoptosis (CC3+), given the larger amount of cell death in the original tumor compared to mouse. The ICC cuboids were loaded into a microtrap device (single array with 96 traps, ∼1 cuboid/trap with flow and manual guidance, see Methods), then processed either immediately after loading (day 0), or after 4 days in culture. For the ICC cuboids, we confirmed the viability of the cuboids by RTG luminescence (**S8 Fig**). The patient cuboids were markedly more heterogeneous than the mouse cuboids, including in the amount of tumor (regions with many nuclei) versus non-tumor tissue (regions with very few nuclei). In contrast with mouse cuboids that showed minimal initial cell death (after elimination of necrotic areas during preparation before slicing), patient cuboids displayed some regions of cell death even at day 0 in CRLM (**Fig 3J**) and in ICC (**Fig 3K**). Many cuboids had regions of very few, well-spaced cells, likely consisting of mostly stromal regions given the tumor-stromal patterns of cellular density by 2D imaging (**Fig 2D**, **S5 Fig**). There was some increase in CC3+ apoptotic cell death at day 4 in the ICC cuboids (**Fig 3K**, bottom row). At day 0 for both CRLM and ICC tumors, the CD45+ immune cells were present variably and in clusters (**Fig 3J,K**), as we had seen by multi-IHC (**Fig 2D**, **S5 Fig**). At day 4 in ICC cuboids, immune cells were still clearly present (**Fig 3K**). Quantitation by Imaris revealed that, unlike in mouse, mean cuboid volume did not decrease (**Fig 3L**). The preservation of a roughly cuboidal shape and similar volume at day 4 suggested minimal shape changes during culture, consistent with 2D brightfield images (**Fig 1**) and histological sections (**Fig 2D**, **S5 Fig**). Immune cell staining did not decrease, consistent with the quantitation of 2D multi-IHC of CRLM cuboids. In sum, 3D immunostaining of patient cuboids confirmed their heterogenous histology and immune cell distribution.

### Cytokine secretion and immune cell activity

Cytokines, including a large family of chemokines, secreted by both tumor and stromal cells, play key roles in the progression, diagnosis, and prognosis of cancer.^64^ Other studies have examined cytokine secretion in relation to immunotherapy for PDEs/PDOTs.^34,65^ For cuboids, we found abundant expression of cytokines in the supernatant of mouse and human cuboids grown in U-bottom 96-well plates with multiple cuboids per well (**Fig 4**). Luminex measurement of Py8119 breast cancer cuboid supernatant revealed baseline expression of 31 of a panel of 32 cytokines/chemokines (**Fig 4A**). Expression of multiple molecules normally secreted by the innate immune system (e.g., G-CSF, IL-6, and TNFα) is consistent with the large presence of macrophages seen in cuboids by immunostaining above. CXCL10 may be produced by monocytes, endothelial cells, and fibroblasts. The low interferon-gamma (IFNγ) and IL-2 signals in **Fig 4A** were consistent with the low presence of T cells observed in Py8119 mouse cuboids by immunostaining and in Py8119 slices by a previous study.^28^ As tumor cells themselves can secrete many of these cytokines, we also tested lipopolysaccharide (LPS) to stimulate the innate immune system directly. LPS application for 24 hrs led to expected increases in inflammatory cytokines, IL-1α and TNFα, and chemokines CCL3, 4, and 5 (**Fig 4B**). These experiments demonstrated both the baseline secretion of many cytokines by immune and/or tumor cells in cultured mouse cuboids, as well as cytokine responses to LPS by immune cells in those cuboids.

**Fig 4.**
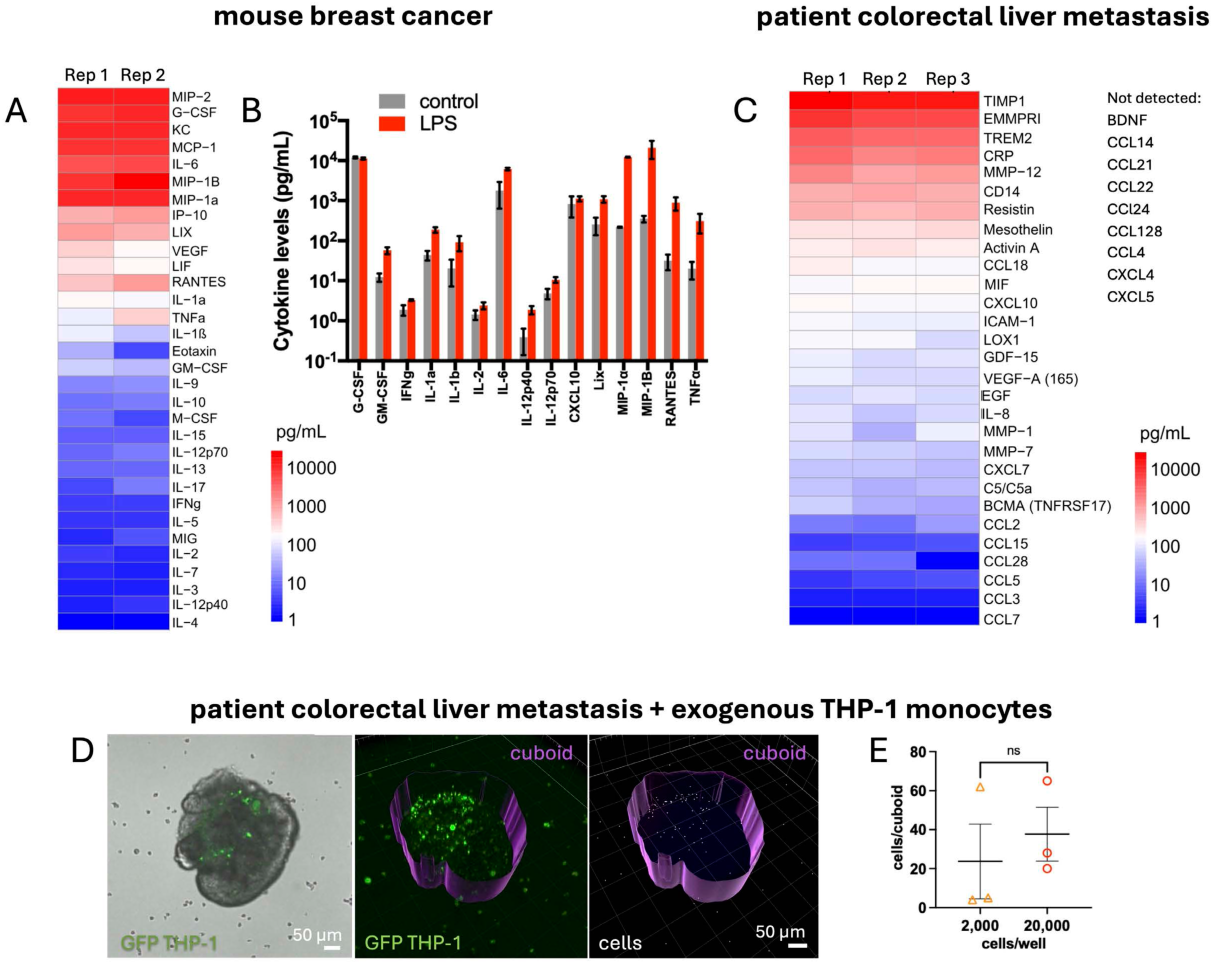
Cytokine secretion and immune cell activity. Cytokine/chemokine secretion in 400-µm cuboids grown in 96 well plates. (**A**) Baseline cytokine secretion from Py8119 mouse breast cancer cuboids by Luminex. 2-6 cuboids/well. Individual replicates (Rep) shown. (**B**) Cytokine response in Py8119 mouse breast cancer cuboids after 24 hour exposure to LPS (lipopolysaccharide, 1 µg/mL) to activate the innate immune system in comparison to control shown in (**A**) (subset of cytokines shown). (**C**) Baseline secretion of cytokines and other factors from human colorectal liver metastasis (CRLM) cuboids. 29 positive signals plus select negative signals of 192 tested by nELISA. 4-6 cuboids per well, day 5. (**D,E**) Patient CRLM cuboids incubated for 4 days with GFP-expressing THP-1 monocytes after 1 day culture from thawing in a 96-well plate. Representative cuboid (20,000 cells/well) with confocal image overlayed on brightfield (left), 3D image of cells in green and surface boundaries of cuboid shadow (middle) and cell identification by Imaris (right). (**E**) Graph of cell number within cuboids. AVE ± SEM. Student T-test.

Cytokines, especially chemokines, have been implicated at all stages of colorectal cancer, including metastasis and recruitment of immune cells.^66^ In supernatant from patient CRLM cuboids (**Fig 4C**), we detected clear expression of 29 cytokines and other secreted molecules of interest at baseline (nELISA on a panel of 192 secreted molecules), most at >10 pg/mL. These molecules included IL-8 (a key pro-inflammatory pro-tumor cytokine^67^), CCL2 (associated with a poor prognosis in CRC),^68^ and CXC chemokines such as CXCL10 (associated with poor survival and prognosis in most human cancers including CRC^64,66^). Other non-cytokine signaling molecules included high levels of soluble Trem2 (Triggering receptor expressed on myeloid cells), a cleaved form of the receptor expressed in TAMs, that is associated with a tumor-suppressive microenvironment.^69^ These cytokines support the use of human cuboids as a model system with expression of cytokines critical to the immunomodulatory crosstalk between tumor and immune cells.

In addition, CRLM cuboids appeared to remain immunologically active, as evidenced by THP-1 monocyte infiltration (**Fig 4D,E**). Single CRLM patient cuboids were cocultured with two different concentrations of monocytes (THP-1 cell line, 2,000 or 20,000 cells per 96-well plate). After 4 days, we localized the green EGFP-expressing monocytes by confocal microscopy (**Fig 4D**). In all of the cuboids, we observed that some of the monocytes had clearly infiltrated into the cuboids (mean of 31 and range 4-65 cells/cuboid), with no statistical difference between the two monocyte concentrations (**Fig 4E**). The infiltration of exogenously added monocytes into cuboids shows that cuboids provide an appropriate substrate for monocyte migration and suggests that the cuboids release an attractive cytokine signal for monocytes. Many of the cytokines identified in **Fig 4C** can attract monocytes, but one, VEGF, was present at a concentration range that could be consistent with attraction near the cuboids.^70^ Furthermore, these preliminary experiments support the potential of cuboids as models with exogenous immune cells (from macrophage to T cells) for cancer immunology studies including immune cell therapies.

### Drug treatment of mouse microtumors

Because of the multi-level heterogeneity of tumor cells and the non-homogeneous distribution of tumor and stromal cells in a tumor, we expected the drug responses to also vary from cuboid to cuboid. To look at single-cuboid variability in drug responses, we treated single Py8119 mouse subcutaneous breast cancer cuboids in a 384-well plate with the cytotoxic chemotherapy drug, cisplatin (CP), the broad-spectrum protein kinase inhibitor staurosporine (STS), or a no-drug control (**Fig 5A-D**). We measured viability by RTG luminescence by IVIS imaging before and after 3-day treatment with drugs. We chose the 3-day timepoint as an appropriate time period for these cytotoxic drugs to produce a measurable change with these viability and cell death assays. At the end of the experiment, we also measured cell death by the terminal SG fluorescent nuclear stain. Both CP and STS resulted in a strong decrease in viability (RTG) and an increase in cell death (SG fluorescence). Despite the relative homogeneity of the mouse Py8119 tumor, there was some variability in the viability before drug treatment and after culture for the control conditions. Using the standard deviation values observed from these drug treatments, power analysis (80% power and α=0.05) estimated that to detect, e.g., a 50% change, we would need ≥2 cuboids for SG and ≥5 for RTG, whereas to detect a 30% change, we would need ≥4 cuboids for SG and ≥12 cuboids for RTG. This difference may arise from the different mechanisms of these assays and would be expected to vary depending on the drug. SG measures loss of membrane integrity while RTG measures metabolic activity proportional to cell number. We expect that the number of cuboids needed for each condition with more heterogeneous tumors, especially with human patient tumors, would be much greater. These experiments demonstrated CP and STS responses at the level of individual Py8119 mouse cuboids using two different assays (viability and cell death).

**Fig 5.**
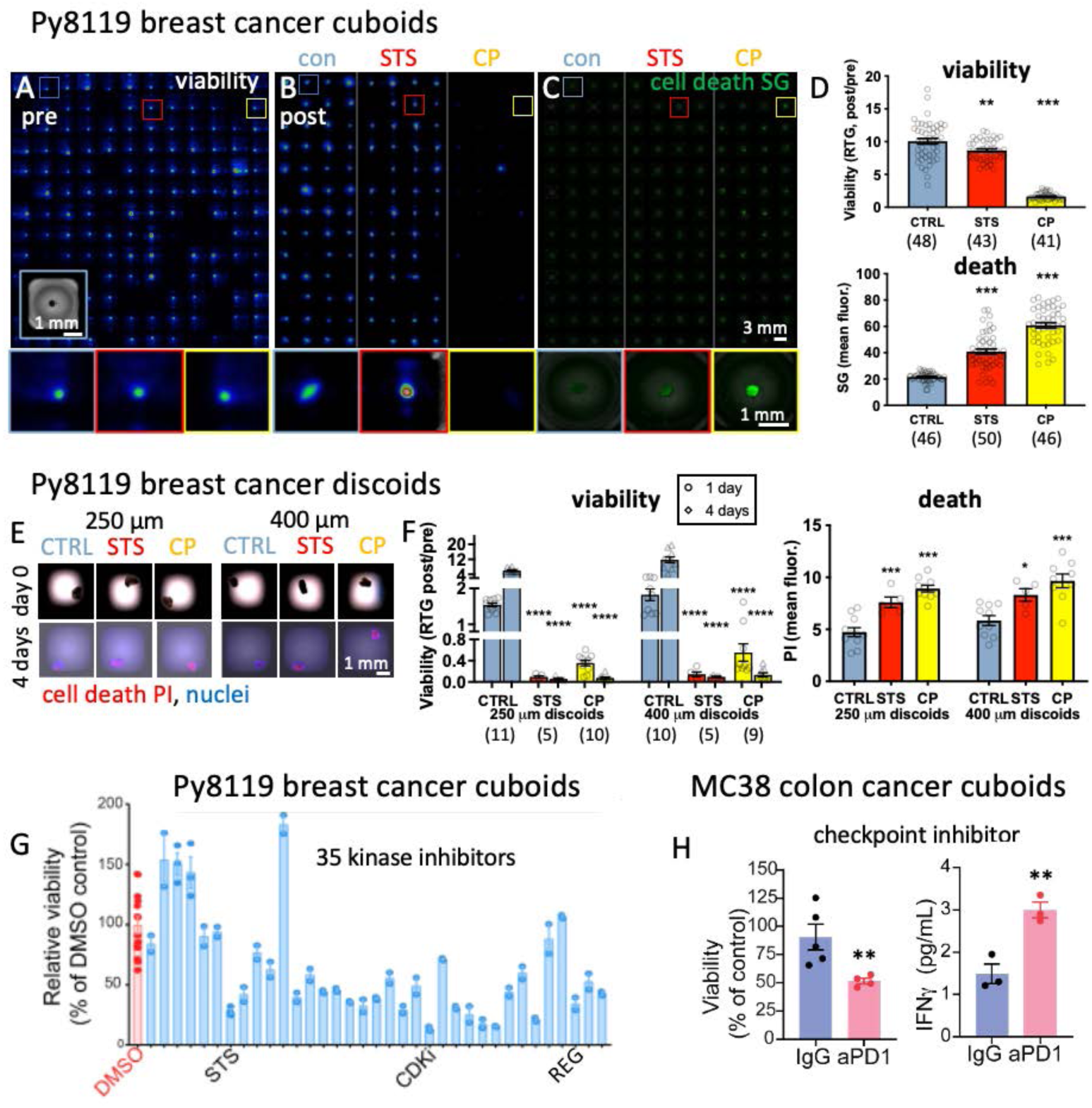
Drug treatment of mouse tumor cuboids and discoids. (**A-D**) Drug responses of individual 400-µm Py8119 mouse subcutaneous breast cancer cuboids before (d2, pre) and after 3 days treatment (d5, post). Manually pipetted with one per well on a 384-well plate, measured for viability by RealTime-Glo (RTG) luminescence by IVIS (**A,B**; pre) and for cell death by SYTOX Green (SG) nuclear stain (**C**). Close-ups shown below. Staurosporine (STS, 2 µM); cisplatin (CP, 100 µM); or control (CTRL). (**D**) Graphs of viability and cell death, with responses of individual cuboids (circles) and AVE ± SEM. One way ANOVA versus CTRL, Dunnet’s post-hoc. **p=0.002, ***p<0.0005. n=41-50. (**E,F**) Drug responses of Py8119 mouse subcutaneous breast cancer discoids. The discoids were cut from an 18G needle biopsy to either 250 µm or 400 µm thickness and robotically placed one per well. Drugs as in (**A**) but applied at d1 (pre). Viability measured by RTG luminescence on a plate reader pre drug at d1, and post drug at d2 and d5 (after 1 day and 4 days). Cell death was measured by red propidium iodide (PI) nuclear stain (on d5) with Hoechst nuclear counterstain (blue). (**E**) Images at d0 and after 4 days of drug treatment. (**F**) Graphs of viability and cell death. AVE ± SEM. One way ANOVA, Dunnett’s post-hoc. *p=0.027, ***p<0.0005, ****p<0.0001. (**G**) Responses of orthotopic mouse Py8119 cuboids to a panel of 35 kinase inhibitors (1 μM). Viability (post/pre) measured by RTG luminescence before and after 3-day treatment, given as a % of the DMSO control. Duplicates of 2-6 cuboids/well pipetted manually. Staurosporine (STS) and CDK4/6 (CDKi) kinase inhibitor positive controls. Regorafenib (REG) VEGFR inhibitor. (**H**) Responses of MC38 subcutaneous mouse colon cancer cuboids to checkpoint inhibitor immunotherapy for 4 days with anti-PD-1 versus IgG control, with decreased viability (RTG) and increased IFNγ secretion. 6-12 cuboids per well pipetted manually. AVE ± SEM. Student’s T-test *p=0.02, **p=0.007.

We also asked whether this pattern of drug responses would be generalizable to Py8119 mouse breast cancer cuboids in microfluidic devices, derived from tumors from either of two locations, subcutaneous or orthotopic (mammary fat pad) (**S9 Fig**). In these experiments, each cuboid trap contained up to 3 cuboids. For both subcutaneous (performed in parallel with the 384-well experiment above) and orthotopic cuboids, like in 384-well plates, CP and STS treatment both led to strong decreases in RTG viability, with CP more dramatically than STS (**Fig 5D**). These experiments demonstrate the ability of cuboids in microfluidic devices to respond to drugs.

Finally, we evaluated whether discoids would show the same pattern of responses to CP and STS (**Fig 5E,F**). We prepared both 250 and 400-µm wide discoids from 18G needle biopsies taken from an excised Py8119 subcutaneous breast cancer tumor. Individual discoids into a 384-well plate were treated with CP and STS for 5 days. Similar to the cuboids, discoids responded strongly to CP and to STS by viability (RTG) on days 1 and 4, and by cell death (PI fluorescence) at the end. These results show that for drug responses, discoids from needle biopsies behave similarly to cuboids, supporting the feasibility of needle biopsies for drug testing.

### Cuboid responses to kinase inhibitors and to checkpoint inhibition

Functional drug testing on patient cuboids could help address the lack of efficacy of targeted therapies on many cancer patients who are candidates by the genetic profile of their tumors. Protein kinase inhibitors (KIs) are a large class of drugs that are often used as targeted therapies for cancer. To demonstrate the suitability of cuboids for drug screening, we tested a panel of 35 KIs on cuboids from an orthotopic mouse Py8119 tumor in a standard 96-well plate format with 2-6 cuboids per well (**Fig 5G**). After measuring the pretreatment viability by RTG luminescence, we removed high and low luminescence outliers to reduce the variability of the starting condition. We measured the drug responses before and after 3 days of drug treatment, relative to the DMSO vehicle control. For 17 of the drugs, we saw >50% reduction in viability compared to DMSO control, and 9 showed ≥75% reduction. For this screen, we used a relatively high concentration per drug (500 nM) at which we would more likely detect a robust response and at which most of the KIs would be expected to inhibit multiple kinases. The high reproducibility of the responses for duplicate wells seen in this KI experiment using multiple cuboids per well contrasts with the variability seen in the CP experiments using single cuboids (**Fig 5D**), both with the same tumor type though different drugs. We attribute the high reproducibility for KI duplicates to both the use of a pool of cuboids per well as well as to the exclusion of wells with unusually high or low pretreatment luminescence. Testing multiple cuboids per well results in signal averaging and minimizes reagent use, which could facilitate higher-throughput drug testing approaches in some situations. On the other hand, examining individual cuboids helps evaluate the heterogeneity of the response and reduces exhaustion of nutrients. Drug responses were similar with fresh and cryopreserved Py8119 cuboids for two select KI’s used in cancer chemotherapy (crenolanib and regorafenib) and the cytotoxic chemotherapy drug cisplatin (4 cuboids/well; **S10 Fig**). These sets of experiments clearly demonstrated responses of Py8119 mouse cuboids to kinase inhibitors, including ones used as targeted chemotherapy, with responses maintained after cryopreservation.

Checkpoint inhibitor (CI) immunotherapy drugs have been previously shown to act in other µDT formats,^26,44,65^ including identification of features predictive of patient response.^34^ As a proof of concept, we applied the PD-1 CI to cuboids from a subcutaneous MC38 mouse colon cancer, which is known to respond to CIs *in vivo*^71^ and in bulk culture of µDTs in collagen^38,44^ (**Fig 5H**). CIs remove the suppression on CD8+ T cells, permitting the T cells to attack and kill the target cancer cells. Given the greater variability in CI responses compared to KI responses, we tested multiple wells for each condition, with 6-12 cuboids per well. After four days exposure to PD-1, we saw both a ∼40% decrease in viability in comparison to IgG control, as well as a ∼2-fold increase in IFNγ, a sign of activation of the immune system. This simple test confirms that cuboids can be used as immunotherapy drug evaluation platforms.

### Drug evaluation with patient cuboids

With the microtumors from patient biopsies we were able to perform drug efficacy tests to evaluate clinically relevant therapies (**Fig 6**). We tested potential treatment options on cuboids from a CRLM patient who had undergone chemotherapy prior to resection, including one cycle with FOLFOX (a combination of folinic acid, 5-fluorouracil or 5-FU, and oxaliplatin; same CRLM as in **Fig 2**). This tumor, like all of the CRLM tumors obtained in this study, was microsatellite-stable (MSS). We tested large numbers of individual cuboids in a 384-well plate either using the cytotoxic components of the standard CRC chemotherapy regimens^72^ FOLFOX or FOLFIRI (5-FU with either oxaliplatin or irinotecan, respectively), staurosporine, DMSO vehicle control, or a no-drug control (**Fig 6A,B**). (Note that for our culture work we did not include folinic acid.) We measured viability (RTG) before (day 2), then after 3 and 5 days of drug exposure. As an additional readout, we measured cell death with PI staining at the end of the experiment after 5 days of drug exposure. We observed significant decreases in viability (RTG) and increases in cell death (PI) for FOLFOX and FOLFIRI (consistent with the patient’s minimal exposure to those drugs), and for the positive cell death control, STS. The small size of the mean reduction in viability for FOLFOX (20%), FOLFIRI (24%), and STS (30%) may in part reflect the heterogeneity of the cuboids, seen not only in the drug responses but in the baseline signal and in the control and DMSO treatments. Power analysis estimated that we would need to test 25 individual cuboids to detect the 24% decrease with FOLFIRI, and 18 cuboids for the 30% decrease with STS (81% power and α=0.05). These experiments show the feasibility of using patient cuboids to evaluate patient chemotherapy drugs. The experiments also demonstrate that, due to the higher heterogeneity of human tumors, there will generally be a requirement for testing a larger number of individual cuboids in total than with the mouse tumors.

**Fig 6.**
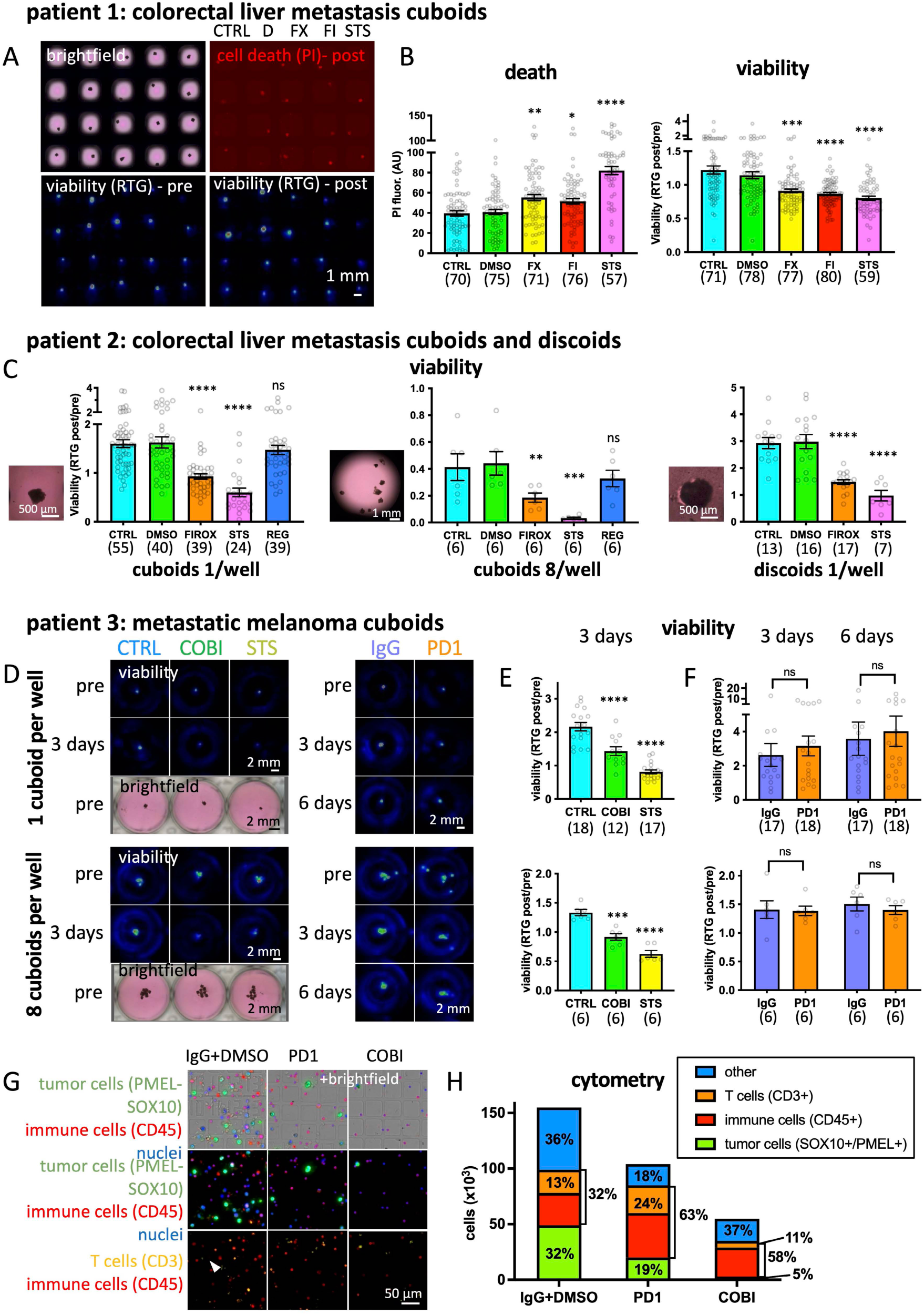
Drug treatment of patient microtumors. (**A,B**) Chemotherapy treatments of individual 400-µm cuboids from a patient colorectal liver metastasis (CRLM). Viability by RealTime-Glo (RTG) luminescence with IVIS measured pretreatment on d2, and after 5 days treatment in a 384-well plate. Death by propidium iodide (PI) at the end. DMSO (0.2%). FOLFOX (FX: 1 µg/mL 5-fluorouracil, 1 µg/mL oxaliplatin). FOLFIRI (FI: 1 µg/mL 5-fluorouracil, 2 µg/mL irinotecan), STS (staurosporine, 1 µM). (**B**) Graphs of viability and death. Responses of individual cuboids (circles) and total number of cuboids in parentheses. AVE ± SEM. PI: One-way ANOVA with Dunnett’s test versus DMSO. *p=0.03, **p=0.002, ***p<0.0001. RTG: Kruskal-Wallis with Dunn’s test versus DMSO. ***p=0.003, ****p<0.0001. (**C**) Chemotherapy treatments of various microtumor formats from a different patient’s CRLM. 400-µm cuboids or discoids were transferred by a robot to 1/well (384-well plate) or 8/well (96 well plate). Graphs of viability of individual wells by RTG on a plate reader pre drug (day 1), then after 6 days of drugs. For discoids, only wells with initial viability over baseline were included. DMSO (0.2%); FOLFIRINOX (FIROX: 1 µg/mL 5-fluorouracil, 2 µg/mL irinotecan, 1 µg/mL oxaliplatin), STS (2 µM), REG (regorafenib, 1 µM). AVE ± SEM. Kruskal-Wallis with Dunn’s (1/well) or One-way Anova with Dunnett’s test (8/well, discoids) versus DMSO. **p=0.009, ***p<0.001, ****p<0.0001, ns = not significant. (**D-G**) Targeted therapy and immunotherapy treatment of 400-µm cuboids from a patient metastatic melanoma lymph node biopsy. Drug treatment starting on d1 with the MEK kinase inhibitor cobimetinib (COBI, 1 µM or media control: CTRL), STS ( 2 µM), or a checkpoint inhibitor (PD-1 or IgG control, 20 µg/mL). (**D**) Representative images of cuboid viability by RTG luminescence (IVIS), with one or eight cuboids per well (96-well plate) pre (d1) or post (d3 or d6) drugs. Inset: color image at d1. (**E,F**) Graphs of viability per well by RTG (post/pre). AVE ± SEM. One-way ANOVA with Dunnett’s test versus control for COBI and STS. Mann-Whitney test for PD1 vs IgG. ***p=0.002, ****p<0.0001. (**G,H**) Cytometry performed on dissociated cells from cuboids after a 3-day drug treatment. Cuboids grown on transwell membranes with 10 cuboids per well, one well per condition. Drugs as above, except for control (0.01% DMSO with IgG). (**G**) Representative images of immunostained dissociated cells, with blue Hoechst nuclear staining. Arrowhead indicates a T cell (CD3 and CD45 positive). (**H**) Graph of the quantitation of cell types as part of the total cell number (% indicated).

We evaluated different microtumor formats on another CRLM tumor, with different numbers of cuboids per well and discoids as well as cuboids (**Fig 6C**). The patient had undergone multiple rounds of treatment, including 3 rounds of FOLFIRINOX (an effective but highly toxic combination of FOLFOX and FOLFIRI used in CRLM) in the past, and most recently FOLFOX. Cuboids (1 or 8 per well) and discoids (1 per well) were exposed to 6 days of FOLFIRINOX. All formats of cuboids and discoids showed significant decreases in RTG viability (luminescence on a plate reader) to FOLFIRINOX (∼40-70%) and to STS, but no response to regorafenib, a KI approved for use in metastatic colorectal cancer. Power analysis for FOLFIRINOX treatments estimated that we would need to test 17 cuboids to detect the 43% decrease with 1 cuboid/well, 11 wells (88 cuboids) for a 58% decrease with 8 cuboids/well, and 4 discoids for the 68% decrease with 1 discoid/well (81% power and α=0.05). In sum, the robustness and flexibility of our approach was supported by the similar responses to three drugs using different formats (cuboid number or discoids) of the same CRLM tumor.

We also examined cuboids from an entirely different type of tumor, a metastatic melanoma (MM), with potential responses to immunotherapy and to targeted therapy. The cuboids were from an inguinal lymph node biopsy of a patient with BRAF-mutant MM (see multi-IHC in **S5 Fig**). The patient’s tumor had progressed after an initial CI response in the past and had recently responded to a MEK inhibitor, binimetinib. We applied CI and KI drugs to the cuboids (either 1 or 8/well) at day 1 for 6 days and measured viability by RTG luminescence from IVIS imaging (**Fig 6D-F**). The treatments were an anti-PD-1 CI, its IgG control, cobimetinib (a clinically approved MEK inhibitor also expected to act on the patient’s BRAF mutation), STS (a multi-KI positive cell death control), or a no-drug control. We saw a clear response to cobimetinib (∼30% decrease) and STS (∼50% decrease for 8 cuboids/well and a ∼65% decrease for single cuboids) by 3 days of treatment (**Fig 6D,E**). The response to cobimetinib is consistent with the tumor’s sensitivity to treatment with a BRAF inhibitor. Power analysis of the cobimetinib drug response for 1 cuboid/well estimated a minimum of 8 cuboids for the ∼65% decrease. For 8 cuboids/well, power analysis estimated a minimum of 2 wells (for a total of 16 cuboids). In contrast, RTG viability measurements did not show any significant decrease for either number of cuboids per well after 3 or 6 days of PD1 treatment (**Fig 6F**). Thus, using a RTG viability assay, we could detect responses of the MM cuboids to a MEK-inhibitor but not to checkpoint inhibition.

Increases in viability from T cell activation and proliferation can mask decreases in viability from tumor cell death, potentially resulting in a lack of viability response such as shown above. To detect cell-specific changes with CKI treatment, in parallel to the above experiment we evaluated cuboid cell numbers by cytometry after treatment with PD1 and cobimetinib (**Fig 6G,H**). Ten cuboids per well were exposed to drugs (PD-1, IgG, or cobimetinib) for 3 days followed by dissociation, immobilization onto PDL-coated substrates with custom grids, and immunostaining for tumor cells (SOX10/PMEL), for immune cells (CD45), and for T cells (CD3) (**Fig 6G**). Cell counts revealed that PD-1 did actually result in reductions in tumor cells and an increase in immune cells and T cells. As expected from the RTG viability results, there was a dramatic reduction in tumor cell number, suggesting that most of the remaining viability in the RTG signal was due to the remaining immune and other non-tumor cells. This result reinforces the importance of matching the type of readout to the type of treatment, sometimes requiring deeper analysis than bulk viability.

### Proteomics of drug-treated mouse cuboids

To provide a deep molecular readout of both drug-induced changes and the state of the TME at once, we adapted a previous “microscaled proteomics” workflow^73^ to the analysis of cuboids (**Fig 7**). We treated cuboids from Py8119 orthotopic mouse breast cancer tumors in 6-well plates for 3 days with several drugs including kinase inhibitors. Treatments included four drugs clinically used for cancer: cisplatin (100 μM), paclitaxel (a cytotoxic chemotherapy microtubule inhibitor; 10 μM), crenolanib (a PDGFR inhibitor; 0.5 μM), and regorafenib (a VEGFR inhibitor; 0.5 μM); as well as doramapimod (a pan p38 MAPK inhibitor used for some autoimmune diseases; 0.5 μM) and a DMSO (0.1%) vehicle control. We prepared samples from replicates of 4 cuboids each, including no treatment (day 4) and pretreatment (day 1) controls. We analyzed the samples by data independent acquisition (DIA) mass spectrometry. With this approach, we consistently identified over 50,000 precursors and 4,000 proteins per sample across treatments using the equivalent of 1 cuboid per injection (**S1 Fig1**). The DIA strategy also afforded high reproducibility (Pearson r>0.95) and precise protein quantifications (median CV<10%) across drug treatments (**S11 Fig**).

**Fig 7.**
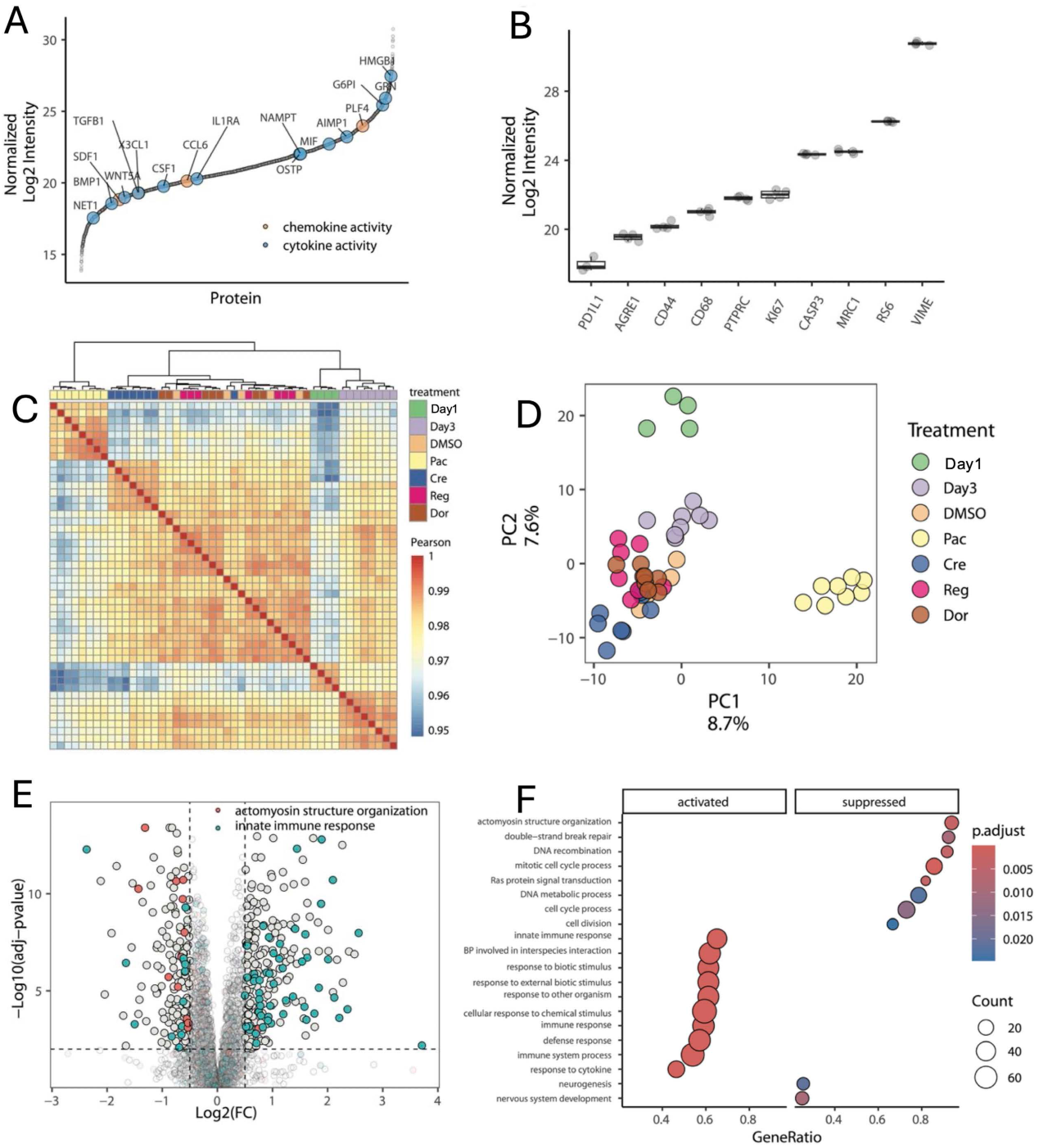
Mass-spectrometry proteomics of mouse breast cancer cuboids and their changes with drug treatments. (**A,B**) Quantification by mass spectrometry of TME proteins from 400-µm PY8119 breast cancer cuboids cultured in 6-well plates at day 1, with 4 cuboids per sample. (**A**) Ranked plot for median normalized log2 protein intensities in day 1 samples; highlighted proteins associated with GO terms: chemokine (yellow) and cytokine activities (blue). (**B**) Box and Dot plot of day 1 normalized protein log2 intensities for observed TME markers.^74^ (**C-F**) Observed changes to the cuboid proteome upon drug treatment from day 1 to day 3. (**C**) Heat map: Pearson correlations for protein abundances across drug treatments and replicates. Day 1, Day3: untreated controls. Treatments (sampled at day 3): **DMSO** (0.1% DMSO), **Pac**: (10 μM paclitaxel), **Cre** (0.5 μM crenolanib), **Reg** (0.5 μM regorafenib), and **Dor** (0.5 μM doramapimod). (**D**) Dot plot: PC1 and PC2 of principal component analysis for protein abundance measurements across treatments and replicates. (**E**) Volcano plot: protein abundance log2 fold changes between DMSO and Paclitaxel treatments on day 3 and adjusted p-values. Log2 fold changes and multiple hypothesis testing adjusted p-values calculated using MSStats linear model. Highlighted proteins in (**grey**) with fold change threshold |Log2FC| > 0.5 and adjusted p-value<0.01; associated with Gene Ontology terms: (**blue**) innate immune response, (**red**) actomyosin structure organization. (**F**) Gene Ontology enrichment analysis of Biological Process terms for upregulated and downregulated proteins between DMSO and Paclitaxel treatments.

Analysis of the expression of select cancer and TME-relevant proteins from day 1 cuboids demonstrated the ability of the proteomics assay to reveal similar information as other assays such as immunostaining, viability, and cytokines, but with multiple types of markers at once (**Fig 7A,B**). Among others, we quantified multiple proteins involved in cytokine and chemokine activities (**Fig 7A**), as well as several TME markers (**Fig 7B**).^74^ These TME markers included proliferation (Ki67) and apoptosis (CC3), key measures of drug response; CD68, a macrophage marker; MRC1 (CD206), a marker of M2 macrophage; PTPRC (CD45) a general marker for immune cells; and VME (vimentin), a marker for tumor cells and the epithelial-mesenchymal transition. Although this approach does not have the cellular resolution of immunostaining or the sensitivity of Luminex cytokine measures, in one test it can evaluate death, proliferation, cell types, and signaling using thousands of target proteins. These proteomic measures could provide a metric to evaluate cuboids for their TME heterogeneity.

We also examined the effects of the different drugs on the cuboid proteome. Measures of similarity of protein abundance by Pearson correlations across replicates (**Fig 7C**) and clustering analysis (**Fig 7D**) revealed that cuboid proteomes undergo substantial and reproducible remodeling in response to drug treatments. As expected, principal component analysis (PCA) to compare the overall similarity in the protein abundance patterns between groups detected a clear difference between day 1 (untreated) and day 3 (treated) cuboids. Paclitaxel, a cytotoxic microtubule inhibitor, seemed to elicit the most dramatic proteome changes in cuboids as reflected by the cluster separation in the PCA (**Fig 7D**, yellow dots). Crenolib also appeared to cause changes in the proteome relative to DMSO, but to a smaller extent. As shown by a “volcano plot” that visualizes all up and down-regulated genes, we identified expected patterns of drug action from paclitaxel treatment, with decreased proteins related to actomyosin structural reorganization and upregulation of proteins related to activation of the innate immune system (**Fig 7E**). With Gene Ontology enrichment analysis (**Fig 7F**), an unbiased method to identify significantly modulated biological processes using the known associations of assigned gene sets, we evaluated the regulated genes between DMSO and Paclitaxel treated samples (**Fig 7F**). Activated processes included multiple immune-related processes and “response to chemical stimulus”. Suppressed processes included double stranded-break repair and Ras signal transduction. This proteomic analysis of small numbers of cuboids demonstrated the potential for rich molecular assessment of cuboids both at baseline and after drug treatment.

### Profiling changes in signaling pathways from drug treatment in mouse cuboids

Assessment of dynamic changes in the activation of protein kinase signaling pathways within cuboids could provide focused, immediate insight into drug action. To complement the unbiased protein analysis by mass spectrometry, which detects thousands of proteins in cuboids, we next focused on identifying post-translational modifications in key growth factor signaling proteins. For this purpose, we used the established reverse phase protein array (RPPA) assay, which relies on tissue extracts and phospho-specific antibodies to detect changes in protein activity.^69^ By applying RPPA, we can accurately identify and quantify post-translational modifications and protein activity changes, even in low-abundance proteins, thus enhancing our understanding of protein dynamics within the TME.

We used reverse-phase protein array (RPPA) to look at dynamic changes in signaling pathways in mouse cuboids with drug treatment (**Fig 8**, **Table S2**). Py8119 breast cancer cuboids were probed with antibodies to look at the activity of 48 proteins, primarily by the phosphorylation state of kinases and kinase-associated proteins, which have critical roles in cancer biology and targeted therapy treatment.^75^ Cuboids were treated for 3 days with cisplatin (10 and 100 μM), doxorubicin (an anthracycline cytotoxic chemotherapy drug, 10 μM), STS (2 μM), or DMSO vehicle control. Two samples of ∼10 cuboids each were probed with 48 antibodies, and 23 yielded high-quality signals. The corresponding 23 proteins revealed distinct protein correlations, functional protein-protein interactions (physical and predicted), and enriched biological processes from gene enrichment analysis. Six proteins of interest showed differences between drug treatments and DMSO (**Fig 8A,B**). Phospho-histone H2A.X (p-H2A.X), a marker of DNA damage, was elevated in some of the drug treatments relative to controls, with loss in high-dose cisplatin and in staurosporine potentially due to excessive cell death lowering the signal. A strong reduction in phosphorylation of ERK, MLC2 (myosin-light chain 2), and BAD (a pro-apoptotic and metabolism regulator) was seen in all drug treatments, consistent with their known involvement with cancer processes (e.g., p-MLC2’s implication in cell proliferation and cytokinesis, tumor progression and the interaction of tumor cells and tumor associated macrophages). Similarly, p-MDM2, a negative regulator of the p53 tumor suppressor, was reduced in the highest concentration of cisplatin and STS only. Other protein signals were unchanged such as LDHA, a monomer of lactate dehydrogenase. Results from the 23 proteins of interest were used to create a Pearson correlation matrix (**Fig 8C**) which measures the similarities of patterns between protein signals with respect to the different treatments. We then created a proposed protein-protein interaction network informed by these experimental responses to drug treatment and published data (**Fig 8D**). This potential insight into the protein signaling networks involved could be followed up with more targeted studies that confirm changes and determine in which cell types they occur.

**Fig 8.**
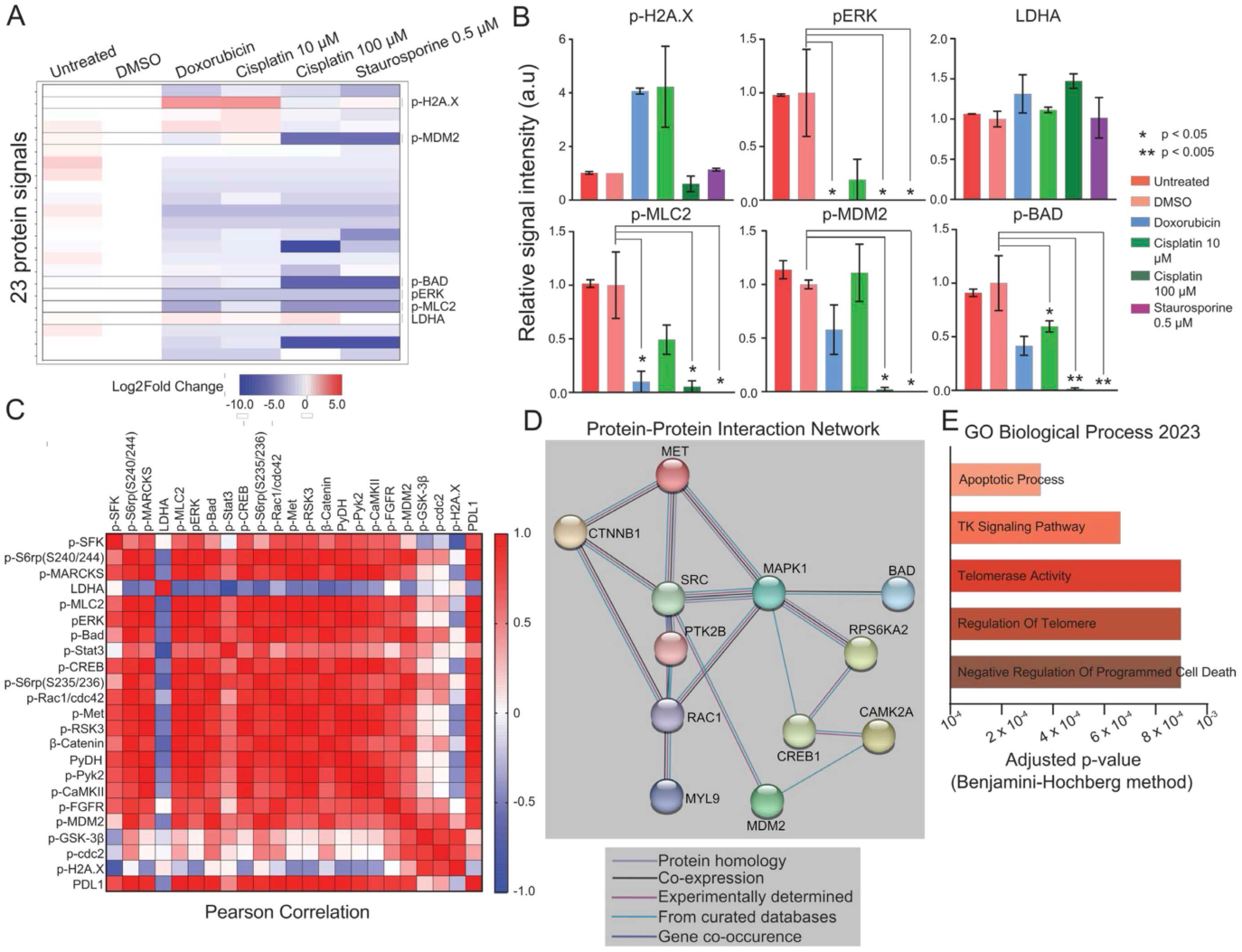
Profiling changes in signaling pathways from drug-treated PY8119 breast cancer cuboids. Reverse-phase protein array (RPPA) was used to profile 48 proteins from PY8119 breast cancer 400-µm cuboid samples after treatment for 3 days in a 6-well plate (untreated, 0.1% DMSO, Doxorubicin 10 μM, Cisplatin 10 μM, Cisplatin 100 μM, or Staurosporine 2 μM). Data from the 23 high-quality signals are shown. (**A**) Heat map representation of relative signal intensities, normalized in a stepwise manner: beta-actin levels, untreated samples, and DMSO treated samples. (**B**) Graphical representation of relative protein levels from annotated proteins in (**A**), normalized in the same process as in (**A**) except without log2 transformation. Except for p-H2A.X DMSO, n=2 per condition and *p<0.05 unpaired t-tests compared to DMSO. (**C**) Pearson Correlation. The correlation matrix depicts the result of a Pearson correlation analysis performed on the 23 proteins of interest using measurements taken from all samples. (**D**) Protein-protein interaction network from positively correlated proteins in (**C**) (StringDB). (**E**) Gene enrichment analysis using the same positively correlated proteins from (**C**) (Enrichr). Adjusted p-values corrected for multiple hypothesis testing by the Benjamini-Hochberg method. Full names of top biological processes: regulation of apoptotic process, transmembrane receptor protein tyrosine kinase signaling pathway, regulation of telomerase activity, and regulation of telomere maintenance via telomerase.

Finally, gene enrichment analysis of the positively correlated proteins (**Fig 8E**) yielded the top biological processes predicted to be involved with these drug treatments. These processes included regulation of apoptosis, transmembrane receptor protein tyrosine kinase signaling pathway, regulation of telomerase activity, and regulation of telomere maintenance. The network of proteins that were substantially downregulated in response to chemotherapy includes receptor tyrosine kinase, such as MET, cytosolic effector proteins like MEK, SRC, FAK, and S6, as well as cytoskeletal proteins such as RAC1. This finding is particularly interesting because Py8119 is a triple-negative breast cancer (TNBC) model known to be driven by the c-MET pathway.^76^ In the majority of TNBC tumors, c-MET is overexpressed, making it a critical driver of cancer progression in these models.^77^ The downregulation of these key proteins in response to chemotherapy suggests a potential mechanism by which the treatment exerts its effects and highlights the importance of the c-MET pathway in the pathology of TNBC. This insight could be valuable for developing targeted therapies aimed at inhibiting the c-MET pathway, potentially improving treatment outcomes for patients with TNBC.

## Discussion

Cuboids offer a cancer microtissue platform whose versatility and ease of interface with automation facilitates their application to studies of drug responses and cancer biology. Their relatively symmetric and regular size facilitates multiple modes of manipulation. Here, we demonstrated the versatility of cuboid manipulation, with manual pipetting of random numbers, manual transfer of individual cuboids, automated selection and transfer by a custom robotic system, and hydrodynamic trapping in a microfluidic device. We also demonstrated the applicability of cuboids to various types of drug testing and their amenability to multiple modes of analysis, including viability measurements, cytokine expression, immunostaining (sections and whole mount), and molecular analysis (proteomics and protein pathway profiling). We also showed (partial) maintenance of drug responses after different cryopreservation protocols that await further optimization for each particular application.

Cuboids benefit from and are limited by the microdissection step much as organoids benefit from and are limited by the expansion step required to grow them. Because organoids are grown, they can be expanded into large amounts of testable tissue (i.e., biobanks) but the growth step also compromises the cellular and physiological similarities between their TME and that of the patient. On the other hand, because cuboids are microdissected, they have an intact TME that preserves the tumor heterogeneity (translating into opportunities for studies of the cancer’s biology) but they are also limited by the biopsy size, which makes biobanking impractical and poses a practical limit for tests performed with the smallest biopsies, such as core needle biopsies. In principle, any solid tumor for which PDEs or PCTS have been published should be microdissectable into cuboids, including breast, colon, prostate, head and neck, and lung patient tumors.^25,26^ The microdissection process could be improved to yield more cuboids per unit volume using sharper, thinner, and/or micromachined^48^ blades. Cuboids and PDEs are presently limited to solid tumors that can be cut and sliced, but the robotic manipulation could be improved to include scooping of soft or ‘mushy’ tumors and their encapsulation into hydrogels. Cuboids that float due to fatty content (e.g., in less differentiated breast cancer or lipid-rich gliomas) could be more difficult to manipulate. This challenge could be addressed by embedding the cuboids in collagen, shown for CRLM cuboids here (**S1 Fig**) and used for microdissected tissues by others.^38,39^

Cuboids cultured for up to one week showed strong viability and cellular diversity, even as they adapted to the acute damage from processing and the limitations of nutrient delivery inherent to tissue culture. While the microdissection process itself inevitably causes cell death on the surface of the cuboid, by the following day we saw no evidence for residual cell death by immunostaining. We chose to focus on the 400-µm cuboids to balance TME representation and diffusion limitations. For mouse cuboids, the central necrosis seen with 400-µm and not 250-µm cuboids suggested that smaller sizes avoid these limitations, and that the larger size can sometimes have a hypoxic core more representative of tumor hypoxic regions. This effect could make cuboids an excellent microscale model to study the effects of hypoxia on tumor development, as previously demonstrated with patient-derived organoids.^78^ Interestingly, the human cuboids studied here did not show this effect, potentially due to less dense tissue. Nevertheless, the majority of both human and mouse cuboids retained multiple cell types by multi-IHC and continued to increase in viability throughout the week.

The preservation of tumor heterogeneity in cuboids provides an opportunity to study the impact of the complex TME on tumor biology and drug responses. Immunostaining clearly revealed the heterogeneity and continued cellular complexity of the TME in cuboids over short-term culture. While our examination of a handful of immune (macrophage, T cells) and stromal (fibroblast, endothelial) cells only covered a subset of the diversity of cell types, we would expect similar representation with many other cell types as seen in other studies of non-passaged, intact tissue models by single-cell RNAseq.^79,80^ Analysis of a larger number of cuboids by single-cell RNAseq in the future could directly assess cell type complexity, with the caveats of poor yield (requiring pooling of cuboids) and sampling bias due to the cell dissociation procedure. More complex assays, such as spatial omics, could provide detailed information on the molecular and cellular heterogeneity between cuboids as they relate to the cuboids’ functional drug responses. The broad range of cytokines observed here from the supernatant of mouse and human cuboids further supports the richness of the tumor and its TME. We observed a slow loss of cells in the mouse cuboids more than in the human cuboids. This decrease in non-tumor cell percentages could result from an increase in the number of other cells (as proliferation was present and viability increased markedly in mouse cuboids), direct cell death (we did not perform double labeling for apoptosis), and/or a loss of cells due to migration from the cuboid. Migration of cells away from the tissue could potentially be mitigated by growth in low attachment plates, or by growth in artificial hydrogels, as suggested by other studies.^53,81^ We did not perform analysis of rarer cell types for individual cuboids given the scarcity of sampling per cuboid and the small size of cuboids. Optimization of size and culture conditions could help to maintain particular cell types as needed, and knowledge of the heterogeneity of those cellular populations between cuboids would guide experiments as well.

As previously observed in PDE cultures,^26^ multiple challenges – and their corresponding opportunities – exist for a better maintenance of the original TME *ex vivo*: mass transport limitations (oxygen, nutrients), preservation of the immune compartment, a suitable external support matrix (e.g., hydrogel-mediated retention of immune and stromal components), and optimization of the culture medium composition.^26^ Future modifications to optimize culture conditions, such as done by Neal et al.^44^ adding IL-2, by Majumder et al.^35^ using patient-derived ECM, or by other groups with hydrogels,^52,53,79,81^ could improve maintenance of cell types. In principle, specialized incubation and perfusion setups (such as macroscale or microscale bioreactors)^26^ could provide more precise control of oxygen levels and metabolism. As the cuboid microscale model presently lacks functional vascular structures, oxygen and nutrient delivery to the cells of each cuboid relies on diffusive mass transport. Recently, the successful vascularization of microscale tumor models by several groups, using a reconstituted human TME^17^ or PDEs,^82^ suggests that a similar approach could be used to vascularize cuboids *de novo*. Even in the absence of circulation, the quantitative reproducibility of diffusive processes at small scales remains an invaluable experimental tool. The microscale size of spheroids, for example, has been used to control and recapitulate hypoxic gradients and necrosis.^78^

As with the intense exploration of models based on organoids and organ-on-chips (OOCs) for tumor development, extravasation and metastasis,^83^ cuboid-based models could potentially be applied to more fundamental questions in cancer biology. Key features of cuboids lend themselves to studies of cancer biology, drug action, and drug delivery. As with other microdissected models, their preservation of the complex 3D tumor TME, with its genetic and cellular diversity and its stromal matrix, enables more relevant studies in culture *ex vivo.*^37,84^ The complexity of the intact tissue models complement other organoid or OOC models whose simpler systems offer higher throughput and greater control of experimental variables, such as the initial tumor’s heterogeneity, an essential factor that could help in studies of tumor biology. The regular sizes of cuboids should improve biological assays by reducing variability and facilitating downstream analysis, such as shown with proteomics and RPPA here. We have also demonstrated that the regular sizes facilitate manipulation by microfluidic traps and robotic handling. Using the same microfluidic device with microtraps and cuboids, our group has shown the ability of flow to counteract loss of endothelial cells in cultured cuboids.^59^ Cuboids should easily combine with more complex OOC models, as done with more traditional cell-derived organoid models.^37^ Here we demonstrated the addition of an exogenous cell type, monocytes, both within and without a collagen hydrogel matrix. Similarly, one could study cellular therapy, such as CAR-T cells, and explore the TME factors that contribute to its failures and success.^37^

Like organoids (PDOs, PDOTS) and PDEs, cuboids are well suited for short-term drug testing studies, particularly due to their complex TME. In our prior study, comparison of drug screening results with RTG found that, on average over three mouse syngeneic tumor models, three times more drugs were effective on 3D cuboid microtumors than on conventional 2D cancer cell lines.^85^ The screening identified a kinase inhibitor, doramapimob, whose effectiveness on microtumors and not on cell lines was due to its action on cancer-associated fibroblasts, and its therapeutic effectiveness was confirmed in 3 mouse models *in vivo*.^85^ Here we demonstrated the suitability of cuboids for drug testing by evaluating the efficacy of cytotoxic chemotherapy, targeted therapy with kinase inhibitors, and checkpoint inhibitors. Supporting the general applicability of cuboid drug testing, our experiments used cuboids from diverse tumor types: two mouse models (breast and colon cancer) and two types of patient tumors (colorectal liver metastases and metastatic melanoma). Cytokine secretion from mouse and human cuboids further supported the functionality of the immune and tumor cells in cuboids. This evaluation confirmed that cuboids could model tumor responses to various treatments, including immunotherapies, much like has been shown by others for other microdissected tumor formats.^34,38,44^ We recognize that tumor responses in patients occur over months and our cuboid cultures only allowed us to look at acute responses to drugs over several days, however, studies similar to ours could be used to provide critical insight into drug action in that particular tumor. Naturally, elucidation of the clinical significance of drug responses in these tests would require further study with clinical correlations. Our study only included pretreated patient tumor biopsies, preventing correlation with future treatment outcomes. Clinical needle biopsies (e.g., with discoids) would be a preferred format for many solid tumors to enable direct pre-treatment testing of drugs for clinical precision oncology.^81^

The semi-automated control of cuboid size and ease of manipulation can facilitate drug testing of microtumors, but requires consideration of how to manage the unique, small µDT pieces. In combination with our low-cost, high-throughput robotic system,^49^ we have the capability to potentially test hundreds of drugs using tissue from a single surgical resection of a patient tumor. Provided the resection is large enough, this approach would facilitate testing drug panels, for example in personalized medicine applications. The higher the heterogeneity of responses, as found in patient tumors, the more cuboids would be needed per drug to determine the overall net effect. The best choice for number of cuboids per well would depend on multiple factors. The higher the variability in the signal (high for CRLM and low for the melanoma), the higher the number of cuboids/wells needed. By measuring the pre-treatment RTG signal, the researcher could assess the variability of the particular tumor and use that information to adjust the sampling per drug condition. Measurement of individual cuboid responses keeps track of their heterogeneity, but at the cost of lower signal-to-noise ratio. For example, with RTG, low signal, non-responsive cuboids were indistinguishable before treatment from viable cuboids with very low signal that would increase in the absence of drug, and that would be extremely sensitive to noise (e.g., all very low signal discoids were removed from analysis for that reason). On the other hand, treatment of cuboids in groups increases sampling, reduces assay complexity and reagent use, increases signal-to-noise with no need to drop wells, and facilitates retrieval for histology or omics analysis. Drawbacks include cuboid-cuboid interactions and increased competition for nutrients. Applications such as immunotherapy which involve unevenly distributed rarer cell types might be better studied with more cuboids per well to ensure representation. In sum, the combination of cuboids and robotics allows flexibility and tailoring of microtissue to the needs of the application.

Our drug response assays were suited to measurement from large numbers of intact, thick tissues. Viability per well (1-8 cuboids) by RTG had the highest throughput, was sensitive to cell death and cell stasis, and was suited for serial measurements. Cell death by fluorescent stains (SG or PI) permitted analysis per cuboid, but was more labor-intensive, susceptible to tissue autofluorescence, and most suited for an endpoint assay. If desired, confocal imaging of the same or similar fluorescent nuclear live/dead stains can give cellular resolution.^6,38^ We showed that these assays can be combined with each other. Both could be followed by molecular assays on the same cuboids, such as protein expression by proteomics, protein pathway profiling (phosphoprotein expression by RPPA), or RNAseq. Overall, a combination of results from mass spectrometry-based proteomics and targeted analysis of signaling events using RPPA performed in parallel can provide a rich analysis of dynamic proteome changes in response to treatments. This integrated approach can yield valuable insights into the molecular mechanisms underlying treatment responses, potentially guiding the development of more effective therapeutic strategies.

Analysis of cell-type specific drug responses within cuboids would require the development of more complex assays than the bulk readouts primarily presented here. Here we used cytometry to reveal changes in cell numbers with checkpoint inhibition. Other cell-type-specific assays options include use of cell-derived tumors from mouse tumor cells stably expressing luciferase, viral transduction of cuboids with luciferase expressed under cell-type specific promoters, or cytokine readouts, such as the IFN-γ secretion shown here. Experiments with dissociated cells, such as flow analysis and single-cell RNAseq, can yield valuable cell type information, but presently require large amounts of tissue per sample due to low yield and suffer from dissociation bias. Confocal live imaging of cuboids could enable the inspection of the interior of live cuboids with fluorescently labeled cells (the limitation of light penetration could be addressed by multi-photon imaging), but new confocal approaches would be needed for higher throughput. Although the deeper insight offered by proteomics and RPPA did not have single-cell resolution, further studies could localize specific target changes to particular cell types, e.g., with immunostaining, single-cell RNAseq, or spatial transcriptomics, as has been explored for other _PDEs.26,40,86_

While more costly and labor-intensive, molecular readouts such as proteomics offer alternatives that could help to maximize the amount of information extracted from drug testing of precious microtumor samples. Over the last decade and largely driven by the efforts of NCI’s Clinical Proteomic Tumor Analysis Consortium (CPTAC), mass spectrometry (MS)-based proteomics has provided profound insights into the molecular mechanisms, diagnosis, prognosis, and treatment of various types of cancer.^87^ Discovery proteomics pipelines can now achieve very high (femtomole) sensitivity for thousands of proteins in low input samples such as individual organoids^88^ or even single cells.^89^ However, the massive efforts by the CPTAC have been undertaken on frozen samples (without culture) and do not provide functional information on drug responses. Recently, the combination of proteomics with microdissection techniques or “microscaled proteomics”^73^ has permitted evaluation of drug responses on sequential core needle biopsies pre- and post-treatment. Proteomics of drug-treated cultured explants (∼1 mm^3^) derived from patient prostate cancers helped reveal some of the underlying molecular and cellular changes underlying the drug response^90^ and suggested the power of application to even smaller tumor pieces such as cuboids.

Further developments of “microscaled proteomics”^73^ for cuboids and other PDEs could enable more sensitive and informative single-cuboid analysis, providing molecular maps of individual cuboids with cell-type information, as well as both unbiased and targeted drug response readouts. Previous proteomic analysis of HSP90 pathways in prostate patient-derived explants used randomly generated larger tissues (mm^3^). Microscaled individual cuboid proteomic analysis could more directly address the expected link between spatial heterogeneity across the tumor and heterogeneous drug responses.^91^ MS-based proteomics is commonly regarded as an expensive technique that is not suitable for multi-well analysis. However, isobaric labeling, e.g., using Tandem Mass Tags (TMTs),^92^ a widely used technique for protein quantification by MS, allows for multiplexing and high throughput. With simultaneous quantification of thousands of proteins across multiple samples in a single analysis, TMT labeling has become the approach of choice by the CPTAC to characterize tumor proteomes,^93^ and could provide a key advance towards scaling up analysis to single cuboids. A complementary development would be the application of targeted proteomics to cuboids (in addition to the unbiased discovery proteomic screen presented here). Targeted proteomics could possibly achieve even greater (attomole) sensitivity and precision quantification of subsets of proteins of interest,^94^ markers of cell composition or immune response^95^ and proteins in pathways relevant for tumor biology.^96^ While immunohistochemical (IHC) imaging provides the ability to localize and detect very rare cells,^97^ proteomics provides superior and unbiased depth and quantitative capabilities, with measurements of hundreds of proteins at once instead of a handful for IHC.^94,98^ Targeted proteomics could be leveraged to find key markers of interest, including drug response markers (for more sensitive or rapid analysis) or cell type markers (for understanding the cellular heterogeneity of the tissue).

Together, this data supports the potential for cuboids as a model system to help bridge the critical gap for cancer biology and cancer drug testing between organoids (which have high throughput and lack TME) and human patients. Cuboids represent a way to optimize and automate the preparation, utilization and manipulation of microdissected tissues (PDEs and PCTSs). Like organoids, cuboids can be integrated with many modalities of assays,^50,99^ allowing researchers help tackle outstanding problems in cancer biology and clinical oncology.

## Materials and Methods

### Cell Culture

The Py8119 syngeneic mouse breast adenocarcinoma cell line (American Type Culture Collection (ATCC), CRL 3278) was grown in DMEM/F12 (for subcutaneous injection) or F-12K (for orthotopic injection) supplemented with 5% fetal bovine serum (FBS) and 1% penicillin/streptomycin. The MC38 syngeneic mouse colon carcinoma cell line (Kerafast, ENH204-FP) was cultured in DMEM supplemented with 10% FBS and 1% penicillin/streptomycin. The THP-1 cells constitutively expressing eGFP/Firefly luciferase under the EF1α promoter (BPS Bioscience, 82148) were grown in RPMI 1640 (with L-glut and HEPES) supplemented with 10% FBS and 1% penicillin/streptomycin. Tissue culture reagents were obtained from GIBCO, ATCC, or Fisher. Cell lines were tested to be mycoplasma-free (Fred Hutch Cancer Center service or MycoStrip, InvivoGen).

### Tumor Generation for Mouse Model

Mice were handled in accordance with institutional guidelines and under protocols approved by the Institutional Animal Care and Use Committees at the University of Washington, Seattle (Protocol 3344-02) or at the Fred Hutchinson Cancer Center (Protocol 50973). Mouse syngeneic tumor cell lines (Py8119 and MC38) were injected into C57BL mice (Jackson Laboratories), >6 week old female (Py8119) or male (MC38) mice. 1-2 x 10^6^ cells were injected subcutaneously (Py8119, MC38) or orthotopically (Py8119, mammary fat pad injection) in 50% Matrigel (Corning) under isoflurane anesthesia. All efforts were made to minimize animal suffering. Tumor size and mouse health were monitored at least 3 times per week. Mice were euthanized using carbon dioxide for tumor harvesting at <2 cm^3^, or earlier if the tumor developed ulceration or if the mouse exhibited any other signs of morbidity as outlined in the University of Washington IACUC Tumor policy. We used 47 animals, of which one died unexpectedly, likely secondary to tumor ulceration. If not used immediately, the tumor was stored up to overnight in Belzer-UW cold storage medium (Bridge-to-Life Ltd).

### Human Tissue

Human tissue was obtained with written informed consent and treated in accordance with Institutional Review Board approved biorepositories at the University of Washington, Seattle. Inclusion criteria were all adult patients with the given cancer type at the University of Washington Medical Center with the exclusion of children. De-identified tumors were freshly procured and processed for slicing the same day as the resection. All of the patient colorectal liver metastases (CRLM) were microsatellite stable (MSS). Patient 1 (ICC5, **Fig 1G,H**) was a 68 year-old female with a history of left renal cell carcinoma one year prior, on pembrolizumab, with incidentally found intrahepatic cholangiocarcinoma. Patient 2 (CRC8, **Fig 1I**, **Fig 1M** orange, **Fig 2**, **Fig 6A,B**) was a 29 year-old female with rectal cancer with recurrent metastasis to the liver, after initial treatment with CAPOX (capecitabine and oxaliplatin) followed by only one cycle of FOLFOX for 2 months. Patient 3 (ICC6; **Fig 1K**, M; **S1 Fig**) was a 54 year-old female with multiple comorbidities and advanced intrahepatic cholangiocarcinoma (FGFR3 gain-of-function variant), stable disease after ∼1 year treatment with durvalumab, cisplatin, and gemcitabine, stopped 4 months before resection. Patient 4 (CRC7, d0 and d2 sizes) was a 54 year-old male with recurrent CRLM, after liver ablation and chemotherapy with cisplatin, irinotecan, leucovorin and 5-fluorouracil. Patient 5 (ICC3, **Fig 1M** dark blue) was a 40 year-old female with biliary ductal adenocarcinoma, resected after neoadjuvant treatment with gemcitabine, cisplatin and Abraxane. Patient 6 (MM1, **Fig 1M**, **S5 Fig**, **Fig 6**) was a 38 year-old female with widely metastatic melanoma that had progressed after initial response to checkpoint inhibition (anti-PD-1 and anti-CTLA-4). The tumor had a VPS54:BRAF fusion and responded to the BRAF inhibitor binimetinib prior to the inguinal node biopsy. Patient 7 (CRC14, **Fig 1M** dark purple, **S1 Fig**, **S6 Fig**, **S7 Fig**) was 65 year-old male with metastatic rectal adenocarcinoma to the liver with primary rectal tumor in situ and previous liver ablation. He had new liver metastases and was on FOLFIRI + Cetuximab. Patient 8 (ICC7, **S1 FigD**) was a 69 year-old male with mixed hepatocellular carcinoma-intrahepatic cholangiocarcinoma and history of hepatitis C. Patient 9 (ICC4, **S5 Fig**) was a 56 year-old male with untreated intrahepatic cholangiocarcinoma. Patient 10 (ICC2, **S8 Fig**, **Fig 3K-M**) was a 70 year-old female with untreated intrahepatic cholangiocarcinoma. Patient 11 (CRC4, **Fig 3H**) was a 42 year-old female with CRLM after neoadjuvant therapy (FOLFOXIRI with cetuximab). Patient 12 (CRC5, **Fig 4I**) was a 66 year-old female with an untreated liver metastasis from cecal adenocarcinoma. Patient 13 was a 43 year-old male (CRC6, **Fig 3J**) with colorectal carcinoma metastatic to the liver, with resection after being off chemotherapy for a year. Patient 14 (CRC15, **Fig 3N,O** & **Fig 6C**) was a 66 year-old male with history of MSS adenocarcinoma of the cecum with liver metastases (diagnosed 11/2024, resected 1/2026), initial resection with neoadjuvant and post resection FOLFIRINOX, a new liver metastasis treated by FOLFOX, which was stopped ∼4 months prior to resection.

### Cuboid Generation and Culture

After removing necrotic regions from the tumors, we generated cuboids as previously described.^8^ For human tumors, and unless otherwise specified for mouse tumors, slices were cut using a Leica VT 1200 S vibrating microtome or MZ5100 vibratome (Lafayette Instruments) after embedding tissue punches (6 mm diameter, Harris Uni-Core) in 1-2% low-melt agarose. Otherwise (experiments with mouse cuboids in microfluidic devices), we cut small pieces of tissue into slices using a tissue chopper (McIlwain) after attaching the tissue to a poly(methyl methacrylate) (PMMA) disk with a thin film of cyanoacrylate glue. We cut slices into cuboids with the tissue chopper, then gently dissociated the cuboids with a transfer pipette. The tissue chopper can be used for all three cuts in sequence (including the first cut) to reduce the duration of the procedure, but at the cost of higher variability in cuboid sizes. To measure the variability attributable to the chopper, we measured the distance between the indentations left by the blade on the PMMA disk with the 400 µm width with an average of 380 ± 13 µm (CV 3.5%, n=16). If using the robot to pick and place cuboids, we used the robot’s machine vision algorithm to select cuboids by size.^49^ For cuboids in serum-free HFH culture medium manipulated by the robot, we got rid of the treated for 20 min, 37 °C in 0.1 mg/mL DNase I (Roche), followed by washing twice in DMEM. Patient melanoma cuboids were individually selected by hand, avoiding fragments. For mouse cuboid experiments not using the robot, e.g., microtrap device or culture experiments, we filtered the cuboids. For 400-µm mouse cuboids, we filtered the suspension through a 750 µm filter (Pluriselect) to remove large fragments followed by filtration through a 300 µm filter (Pluriselect) to remove small debris. To remove potential contamination introduced during the chopping procedure (performed outside the hood for mouse cuboids) we washed the cuboids at least twice with sterile PBS or DF12 either in a tube or a 100 µm cell strainer (Corning or Falcon).

Cuboids and discoids were cultured in 6 well plates (<50 cuboids per well, 3 mL per well), 96-well plates (200-250 μL), or 384-well plates (120 μL). For cuboids and discoids from patient, MC38, and select Py8119 tumors (cytokine, checkpoint inhibitor, and drug panel experiments, unless otherwise specified), the culture medium was HFH: Williams’ Media E (Sigma) supplemented with nicotinamide (12 mM), L-ascorbic acid 2-phosphate (50 mg/mL), D-(+)-glucose (5 mg/mL) from Sigma; sodium bicarbonate (2.5%), HEPES (20 mM), sodium pyruvate (1 mM), Glutamax (1%), and penicillin–streptomycin (0.4%) from Gibco; and ITS + Premix (1%) and human EGF (20 ng/mL) from BD Biosciences or Corning. For other Py8119 experiments (e.g., drug testing immunostaining, RPPA, and proteomics), cuboid culture medium was DMEM/F12 with 5% heat-inactivated FBS and 0.1% penicillin-streptomycin. We prepared 80% collagen (Corning rat tail collagen type 1, 354236, 3-4 mg/mL) with 10% 10x PBS, and 10% serum-free medium, neutralized with filtered-sterilized 1M NaOH to pH ∼7.2. Cryopreservation was performed according to manufacturers’ instructions using Recovery (GIBCO), CryoStor CS10 (STEMCELL Technologies), or CRYO-SOL (Organotech), with less than 100 cuboids per 500 mL solution.

For patient cuboid and monocyte coculture experiments, patient-derived colorectal cancer liver metastasis cuboids previously cryopreserved in CryoStor were thawed and allowed to recover for 24 h at 37 °C and 5% CO₂. Individual cuboids were then co-cultured in 200 µL of HFH medium supplemented with heat-inactivated 5% FBS with THP-1 cells constitutively expressing eGFP under the control of the EF1α promoter, with either 20,000 or 2,000 cells per well in a 96-well plate. Co-cultures were maintained at 37 °C and 5% CO₂ for 4 days, after which cuboids were transferred to a glass-bottom 96-well plate for confocal imaging.

### Drug Treatment and Functional Assays

To measure viability, RealTime-Glo (RTG) (Promega) was added at 1x or 0.5x and after overnight incubation, the baseline luminescence was read by IVIS (PerkinElmer). Drug was added to the well, and luminescence was read again after incubation with drug without further addition of RTG. Low viability for patient cuboids (individual or wells) was defined as less than 33% RTG luminescence of the mean (excluding the high outliers), except for in collagen (<10% of mean). For fluorescent nuclear cell death endpoint assays, we added SYTOX Green (1/50,000; Invitrogen) or propidium iodide (1 µg/mL), with or without Hoechst (16 µM or 1x NucBlue, Invitrogen), incubated 1 hr at 37 °C, then imaged with or without washing twice in PBS. For the kinase inhibitor drug panel experiments, multiple cuboids were randomly distributed by manual pipetting, and baseline viability was used to remove outliers before randomly assignment of treatments. Cytokine measurements on supernatant were performed by Luminex (Py8119) or by nELISA (CRLM, Nomic Bio).

Cuboids were treated with lipopolysaccharide (Sigma L4516) or drugs. Antibodies to checkpoint inhibitors (10 μg/mL) were mouse anti-human PD-1 antibody (BD Biosciences, 562138, clone EH12.1), mouse IgG1 control (BD Biosciences 554721), rat anti-mouse PD-1 antibody (Leinco Technologies, P372), and rat IgG2a control (Leinco Technologies, R1367). Drugs for treatment of cuboids included cisplatin (MedChemExpress), staurosporine (Thermo Scientific), and fluorouracil, oxaliplatin, irinotecan, and cobimetinib (Selleck).

The 35 drug panel for treatment of Py8119 cuboids, in order, was DMSO, Baricitinib (LY3009104, INCB028050), Doramapimod, JNK Inhibitor VIII, TCS ERK 11e (ERK Inhibitor VIII, VX-11e), Rho Kinase Inhibitor III Rockout, PP121, GZD824, Staurosporine, AZD3463, NVP-BVU972, Y-39983 dihydrochloride, cobimetinib, WZ3146, Aminopurvalanol A, SB202190 (FHPI), PD 166285 dihydrochloride, BIIB-057 (PRT062607, P505-15), Akt Inhibitor VIII Isozyme Selective Akti-1/2, CP 547,632, Bosutinib, AT7867, Cdk1/2 Inhibitor III, PDK1/Akt/Flt Dual Pathway Inhibitor, SB 218078, PF-3644022, LY 333531 mesylate (Ruboxistaurin), Cot inhibitor-2, PD 173955 Analogue 1, Vargatef (Nintedanib), BIBF 1120), K252a, Nilotinib, GSK650, Regorafenib, Pazopanib, and At9283. Drugs were obtained from MedChemExpress, Selleck Chemicals, TOCRIS, Cayman, or Sigma.

### Microfluidic Device Fabrication and Use

Microchannels were made in a process adapted from Horowitz et al.^8^ with a design similar to the more complex, valved design made by CNC milling in Lockhart et al.^11^ We designed the devices in AutoCAD (Autodesk) and then fabricated them by laser micromachining (VLS3.60, Scottsdale, USA) of PMMA substrates followed by thermal solvent bonding and adhesive bonding. The device has four layers. The 800 µm-thick PMMA channel network layer (Clarex, Astra Products) has laser-cut microchannels that connect the outflow channel(s) to microtraps via binaries. The microtraps were approximately 560 µm in diameter at the top and 620 µm at the bottom. The microchannels were approximately 410 µm wide except for near the traps where they were approximately 320 µm wide. After sonication of the microchannel layer in isopropanol for 1 min, we bonded it to a 300 µm-thick PMMA sealing layer (Clarex, Astra Products) using thermal solvent bonding; we exposed both layers to chloroform vapor (∼3 mm above the liquid) for 4 min, manually pressed them together, then sealed in a heat press for 5 min at 140 °C, 240-260 psi. The well plate layer was laser-cut from 6.35 mm-thick PMMA (1227T569, McMaster-Carr, Elmhurst, IL) and bonded to the channel layer using double-sided 3M™ High-Strength Acrylic Adhesive 300LSE for 2 min at 250 psi in the thermal press. Silicone tubing was bonded to the outlets using cyanoacrylate glue. Optionally, an additional well plate layer was hand-bonded onto the device after cuboid loading using the 300 LSE. The devices were sterilized by running 70% ethanol through the channels.

To load cuboids into the devices, we used negative pressure to pull the cuboids into the traps. Cuboids were suspended in ice-cold 20% polyethylene glycol (8,000 MW, 8k-PEG P2139, Sigma-Aldrich), whose increased density facilitates movement of the cuboids without damaging their viability.^8^ After pipetting the cuboid solution into the well, we applied suction with a 60-cc syringe and a syringe pump (Chemyx Inc.) at 200 mL/hr for 96-microtrap wells or 100 mL/hr for 48-microtrap wells. Mouse cuboids were pushed around the well slowly using either a custom window (single-well, 96-trap device), or a comb (3-well device). For the scarce human cuboids, under visualization with a VHX-970F microscope (Keyence), cuboids were pushed manually to the vicinity of the traps using forceps to encourage only one cuboid per trap while applying suction. More PEG solution and more cuboids were loaded as needed. The loading process took about 10 min for the 96-trap wells and 1 min for each 48-trap well. Immediately after loading, the solution was exchanged for culture medium. After disconnecting the syringe pump, the device was placed into a 10 cm petri dish for culture.

### Immunostaining and histology of tissue sections and whole mount tissues

For multi-immunohistochemistry experiments, tissue pieces and cuboids were fixed in 4% paraformaldehyde and washed with PBS. Cuboids were then embedded in Histogel (Epredia) plugs. Tissue was then processed for microarrays, paraffin sectioning, hematoxylin & eosin (H&E) staining, and multi-immunohistochemistry (multi-IHC) by the Experimental Histopathology Service at the Fred Hutchinson Cancer Research Center. Paraffin section (4 µm) were stained using the Leica BOND Rx stainer and processed with either two 5-plex (mouse) or three 3-plex (human) stains. Lists of antibodies are given in **Table S1**. Staining for tumor cells (CK19) was weak and variable so was not used for quantitation. DAPI was used as a nuclear counterstain.

For peroxidase immunostaining (**S1 Fig**), tissue was fixed in 4% paraformaldehyde overnight then cryoprotected with 30% sucrose/PBS overnight two times. Cryosections (14 µm thickness) were then processed for H&E or for immunostaining. For immunostaining, we pretreated with 0.6% hydrogen peroxide in methanol for 30 min, washed, and then performed antigen retrieval by steaming for 30 min in 10 mM sodium citrate, 0.05% Tween 20 (Sigma), pH 6.0. After at least 30-min block in blocking solution (Tris-NaCl-blocking buffer or TNB buffer, Perkin Elmer, with 0.1% Triton X-100), we incubated the tissues with rabbit primary antibodies (diluted in TNB) overnight at 4 °C. Antibodies were: active cleaved caspase 3 (1/600, Cell Signaling), Ki-67 (1/1,000, AbCAM, ab15580), CD31 (1/200, AbCAM ab28364), or CD45 (1/1,000, AbCAM, ab10558). After incubation of the tissues with an anti-rabbit peroxidase polymer (Vector Labs MP7401) for 30 min, we visualized the staining with 3,3’-Diaminobenzidine (DAB, Vector Labs) and a light hematoxylin counterstain. Washes between steps were done with PBS except for just before DAB incubation, done with 10 mM Tris pH 7.5.

We performed whole-mount immunostaining in the microfluidic devices with a protocol adapted from Li et al.^100^ Cuboids were incubated in block (PBS + 0.3% Triton + 1% BSA + 1% normal mouse serum) for >12 hrs at 37 °C on a shaker, then in primary antibody diluted in block for 2 days at 37 °C on a shaker. Primary antibodies were Alexa 647 anti-mouse CD45 (1:200; Biolegend), Alexa 488 anti-mouse CD31 (1:250; Biolegend), Alexa 647 anti-human CD45 (1:400; HI30 Biolegend 304056), rabbit anti-human/mouse cleaved-caspase 3 (1:300; Cell Signaling, 9661), followed by 1:500 546-Donkey/Goat anti-rabbit (or other). We then washed the cuboids 3 times with PBS at least 8 hrs total at 37 °C on a shaker. For the unlabeled primary we incubated with Alexa 546 donkey anti-rabbit IgG (1:500; Invitrogen) diluted in block for 2 days at 37 °C on a shaker followed by washing as above. DAPI 1 µg/mL was added to the last antibody step. We then cleared the tissue by incubation in previously prepared Ce3D solution overnight at room temperature with shaking. The day 0 human ICC cuboids were immunostained in a centrifuge tube and imaged in a custom well.

Whole-mount pseudo-H&E staining (TO-PRO-3 and eosin, ethyl cinnamate clearing) and open-top light-sheet microscopy of a needle biopsy and cuboids (embedded in low-melt agarose) were performed by Alpenglow Biosciences, Inc., with methods described previously.^8^

### Imaging and Image Analysis

We took images (except for cytometry, see below) using a Canon DS126601 on a Nikon SMZ1000 dissecting scope, a Nikon Eclipse Ti microscope, a Keyence BZ-X800 microscope, a Keyence VHX microscope, an Olympus Fluoview 3000 confocal microscope (monocyte experiments), or a Leica SP8 confocal microscope (10x, 2 µm Z steps, at the Lynn & Mike Garvey Imaging Core at the University of Washington). For confocal imaging of 3D immunostaining, we only imaged the cuboids at the bottom of a well. For general image manipulation we used FIJI. We used FIJI to estimate cuboid size and for measurements of mean fluorescence. We defined regions of interest using a binary mask and removed clumps of cuboids from analysis (in bulk culture). For RTG luminescence taken from IVIS images, we defined wells as regions of interest using a black and white photograph taken in parallel.

Multi-IHC imaging utilized the Vectra Polaris Quantitative Pathology Imaging System (Akoya Biosciences, Marlborough, MA). Images were spectrally unmixed using Phenoptics inForm software and exported as multi-image TIF files. For multi-IHC image analysis, we used HALO and HALO Link (Indica Labs). For cellular analyses, cells were identified by nuclear DAPI stain, then cytoplasmic boundaries were drawn. For antibodies with clear staining, we applied the same settings to all tissues with similar staining, applying minor adjustments as required to staining groups with varied overall signal (Ki-67, CD45, CD3, CD8, and F4/80). For CD3+ determination, we included all cells with CD8 or CD3 staining as CD8 could occlude CD3. For NKp46 staining, which had few cells in cuboids in days 1 through 7 with high background, we performed an automated count for the initial tumor and did a manual count for the cuboids. For area analyses (CD31), we applied the same settings to all tissues of that tumor type. We used the background staining from CK19 or from NKp46 to determine the total tissue area.

For 3D image analysis, we used Imaris (Oxford Instruments) through the Lynn & Mike Garvey Imaging Core. Cuboid volume was determined by creating a surface with the DAPI channel or background from another channel. CD31 and CD45 surfaces were created using Labkit for machine learning. For CD31, we applied a gaussian blur before further analysis. We trained the classifiers for days 0 and 2 separately on a stack of two cuboids initially, then applied the classifier to all the cuboids in that day’s group. For CD45 on mouse, we trained the classifier on day 0, then applied those settings to all the cuboids.

Analysis of monocytes and cuboids was performed using Imaris. Monocyte identification and quantification within cuboids was performed using the spot function, with machine learning to create a setting applied to all cuboids. Cuboid boundaries were defined using the surface function to create a columnar space defined by the brightfield shadow. The few remaining extracuboidal monocytes after transfer from the culture well were easily distinguished.

### Tissue cuboids treatments and immunofluorescence staining for cytometry

Human melanoma cuboids were cultured in HFH. Following 72-hr treatment with drugs (20 μg/mL anti-PD1 or IgG; 1 μM Cobimetinib), cuboids were dissociated using the human tumor dissociation kit (Miltenyi Biotec). Specifically, 50 μL Enzyme H, 25 μL Enzyme R, and 6.25 μL Enzyme A were added to 1.1 mL serum-free RPMI 1640 media (ThermoFisher) to prepare the dissociation solution. Cuboids were rinsed once with PBS, transferred to the dissociation solution, and incubated at 37 °C with 1800 rpm shaking for 15 min. The resulting cell suspensions were filtered through a 30 μm strainer (Miltenyi Biotec) to remove undigested tissue. For staining, cells were seeded onto poly-L-lysine (Gibco)-coated micropatterned PDMS (polydimethylsiloxane, SYLGARD 4019862, Corning) substrates on glass slides. Micropatterned substrates were fabricated by standard soft lithography techniques.^101^ Cells were fixed with 4% paraformaldehyde (Santa Cruz) for 15 min at room temperature. Cells were then permeabilized with 0.5% Triton X-100 (Sigma Aldrich) for 15 min and blocked with 5% BSA (Sigma Aldrich) for 60 min at room temperature. Subsequently, cells were stained overnight at 4 °C with Ki67-FITC (Invitrogen, 11-5698-82); SOX10, PMEL (Atlas Antibodies, HPA001758, HPA031649); and CD3-APC (Biolegend, 300458) in PBS containing 1% BSA. Secondary antibody staining was performed for 60 min at room temperature using anti-rabbit IgG (H+L) secondary antibody, TRITC conjugate (Jackson ImmunoResearch, 711-025-152), in the presence of 0.2% BSA. Finally, cells were labeled with Hoechst 33342 (BD Biosciences) for 10 min at room temperature and imaged using an ImageXpress Micro microscope (Molecular Devices). For quantitative analysis, the number of tumor cells and T cells was quantified in ImageJ by overlaying the positive cell type segmentation (generated with the ImageJ plugin ilastik) with the corresponding positive nuclear signals.

### Cuboid Sample Preparation for Mass Spectrometry Analysis

Four PBS-washed cuboids per replicate were resuspended in 40 μL of lysis buffer (6M GuHCL, 50 mM Tris pH 8.2, 75 mM NaCl, 10 mM TCEP, 30 mM CAA). The cuboid suspension was subjected to 4 rounds of agitation at 90 °C for 5 min followed by 15 min of bath sonication at 40 °C. Alkylation was quenched with 15 mM DTT for 30 min at room temperature and the samples were diluted 5-fold with Tris 50 mM pH 8.9 prior to digestion, which was carried out overnight at 37 °C using LysC endoproteinase at a 5 ng/μL concentration. Digestion was quenched with 1% TFA prior to desalting with oasis prime HLB μElution (Waters). Desalted peptides were dried down by vacuum centrifugation.

### Mass Spectrometry Data Acquisition

Lyophilized peptide samples were resuspended in 3% ACN, 5% formic acid and subjected to liquid chromatography coupled to tandem mass spectrometry (LC-MS/MS). Samples were loaded into a 100 μm ID x 3 cm precolumn packed with Reprosil C18 1.9 μm, 120 Å particles (Dr. Maisch). Peptides were eluted over a 100 μm inner diam. (I.D.) x 30 cm analytical column packed with the same material housed in a column heater set to 50 °C and separated by gradient elution of 6 to 38% B (A: 0.15% FA, B: ACN 80% and 0.15% FA) over 65 min and 38% to 50% B over 8 min at 400 nL/min delivered by an Easy1200 nLC system (Thermo Fisher Scientific) with a total 90 min method length. Peptides were online analyzed on an Orbitrap Eclipse Tribrid mass spectrometer (Thermo Fisher Scientific). Mass spectra were collected using a data-independent acquisition method. For each cycle a full MS scan (358-1100 m/z, resolution 60,000, and Standard 100% normalized AGC target in automatic maximum injection time mode) followed by 30 consecutive non-overlapping MS/MS scans with 24 m/z wide isolation windows. MS/MS scans were acquired at 30,000 resolution in centroid mode, using HCD fragmentation with 33% normalized collision energy and an 800% normalized AGC target with automatic maximum injection time mode. In alternating cycles, the mass target list for MS/MS scans was offset by 12 m/z to produce DIA windows with half overlap spanning the 363-1095 m/z mass range. Information on Mass spectrometry Data Analysis is supplied in Suppl. Info. The mass spectrometry proteomics data was deposited to the ProteomeXchange Consortium via the PRIDE partner repository (https://www.ebi.ac.uk/pride/).

### Reverse Phase Protein Arrays (RPPA)

Protein microarrays were printed and processed as previously described.^102^ Briefly, cuboid lysates (duplicates of 10 cuboids/sample) from the PY8119 syngeneic mouse breast adenocarcinoma cell line were prepared in 2% SDS lysis buffer (stock recipe, pH = 6.8: Tris⋅HCl 50 mM, 2% SDS, 5% glycerol, EDTA 5 mM, and NaF 1 mM). Immediately before use, we added protease inhibitors and phosphatase inhibitors (β-GP 10mM, PMSF 1 mM, Na_3_VO_4_ 1 mM, and DTT 1 mM) and then printed onto RPPA slides, each containing 16 nitrocellulose pads (Grace Biolabs, GBL505116), using Aushon 2470 microarrayer (Aushon BioSystems). Samples were printed in duplicate and stored at −20 °C until processing. Slides were then washed with 1 M Tris⋅HCl (pH 9.0) for 2 to 3 days to remove SDS and PBS solution for 10 to 15 min, centrifuged with Thermo Sorvall Legend XTR Refrigerated Centrifuge (Thermo Scientific), and scanned with InnoScan 910 AL (Innopsys) to ensure no residual SDS. Slides were washed with PBS solution 2 to 3 times for 5 min each and then blocked with Odyssey blocking buffer (OBB) (Licor, 927–40000) for 1 hr at room temperature. Afterwards, slides were fitted to hybridization chamber cassettes (Gel Company, AHF16), incubated with primary antibodies on a shaker overnight at 4 °C. The following day, slides were washed with PBS thrice for 10 min each, blocked again with Odyssey blocking buffer (OBB) (Licor, 927–40000), and incubated with IRDye-labeled secondary antibodies for 1 hr at room temperature. After incubation, slides were washed in PBS 3 times for 10 min each, centrifuged again with Thermo Sorvall Legend XTR Refrigerated Centrifuge (Thermo Scientific), and scanned using Licor Odyssey CLx Scanner (LiCOR). Total signal intensity was normalized to total beta-actin (Sigma, Catalogue No A1978) and quantified using Array-Pro analyzer software package (Media Cybernetics, Maryland, USA). To enable the generation of the heatmap and Pearson correlation figures, we replaced 0 values with the lowest detected values from the other treatments.

### Statistical analysis

Prism 7 or 10 (GraphPad) were used for graphing and statistics unless otherwise noted. Details on statistical tests used are described for each use in figure legends. In general, we used non-parametric tests instead of parametric tests when data clearly had non-normal distribution (evaluated by D’Agostino & Pearson, Anderson-Darling, Shapiro-Wilk, and Komogorov-Smirnov tests on Prism). For comparison of two samples, we used the student’s unpaired two-tailed T test for parametric data and the Mann-Whitney test when nonparametric. For multiple comparisons, when parametric we used one-way ANOVA with Tukey’s post-hoc or with Dunnett’s post-hoc if comparison to one condition. When nonparametric, we used Kruskal-Wallis with Dunn’s post-hoc test with comparison to a single condition). Post-hoc power analysis (80% power and α=0.05) was performed using https://clincalc.com/stats/samplesize.aspx.

## Availability of data and materials

Multi-IHC and 3D imaging data and analysis are available on reasonable request. The mass spectrometry proteomics data have been deposited to the ProteomeXchange Consortium via the PRIDE partner repository (https://www.ebi.ac.uk/pride/). All other data are available in the main text or in the supplementary materials.

## Acknowledgments

We thank Heidi Kenerson for preparation of human slices. This research was supported by the Experimental Histopathology Shared Resource of the Fred Hutch/University of Washington/Seattle Children’s Cancer Consortium (P30 CA015704).

## Author contributions

Conceptualization LFH, RR, AL, EW, WW, JV, RSY, TSG, AF

Formal analysis LH, RR, MY

Supervision EW, WW, RSY, JV, TSG, AF

Funding acquisition LH, RR, EW, WW, JV, RSY, TSG, AF

Investigation LFH, RR, SZ, YT, AL, NRG, IS, MY, TNHN, EJL, CA, TSG

Visualization LFH, RR, MC, SZ, YT, AL, NRG, IS, CS, MY

Methodology LFH, RR, AL, JV, TSG, AF

Writing-original draft LFH, RR, JV, TSG, AF

Writing-review and editing LFH, RR, JV, TSG, AF

