## Supplemental materials for "Functional evaluation of microdissected tumor ‘cuboids’: a microscale *ex vivo* cancer model that retains a complex tumor microenvironment"

Lisa F. Horowitz* *et al.*

**This PDF file includes:**

Supplemental methods: Mass spectrometry Data Analysis
S1 to S11 Figs
S1 & S2 Tables
S1-S4 Movie captions

**Other Supplementary Materials for this manuscript include the following:**
S1-S4 Movies

**Supplementary methods: Mass spectrometry Data Analysis**

Raw files were converted to mzML format and demultiplexed using MSConvert (https://proteowizard.sourceforge.io/tools/msconvert.html) followed by analysis with FragPipe (https://fragpipe.nesvilab.org/, version 19.1) proteomics pipeline. Searches were performed with MSFragger (https://msfragger.nesvilab.org/, version 3.7) using a target/decoy *Mus Musculus* database (Uniprot fasta proteome: UP000000589_2023-03-02). Search parameters were set to 20 ppm for both precursor and fragment tolerances. LysC was selected as the digestive enzyme with a maximum of 1 missed cleavage and fixed constant carbamidomethylation modification of cysteines (+57.0215 Da) and variable modifications of methionine oxidation (+15.9949 Da) and protein N-terminal acetylation (+42.0106 Da). Search results were post-processed with Philosopher (https://philosopher.nesvilab.org/, version 4.8.1) using Percolator (http://percolator.ms/, version 3.05) to filter the results to 1% False Discovery Rate (FDR) and ProteinProphet (https://proteinprophet.sourceforge.net/) for protein inference. EasyPQP (https://github.com/grosenberger/easypqp) was used to generate a peptide spectral library and quantification was carried out using DIA-NN (https://github.com/vdemichev/DiaNN, version 1.8.2).

Bioinformatic analysis was performed using R (https://www.r-project.org/) and the R Studio environment. MSStats (https://github.com/Vitek-Lab/MSstats, version 4.01) was used to parse DIA-NN quantification outputs, generate and normalize protein quantification and carry out statistical significance analysis. Gene ontology enrichment analysis was carried out using ClusterProfiler.

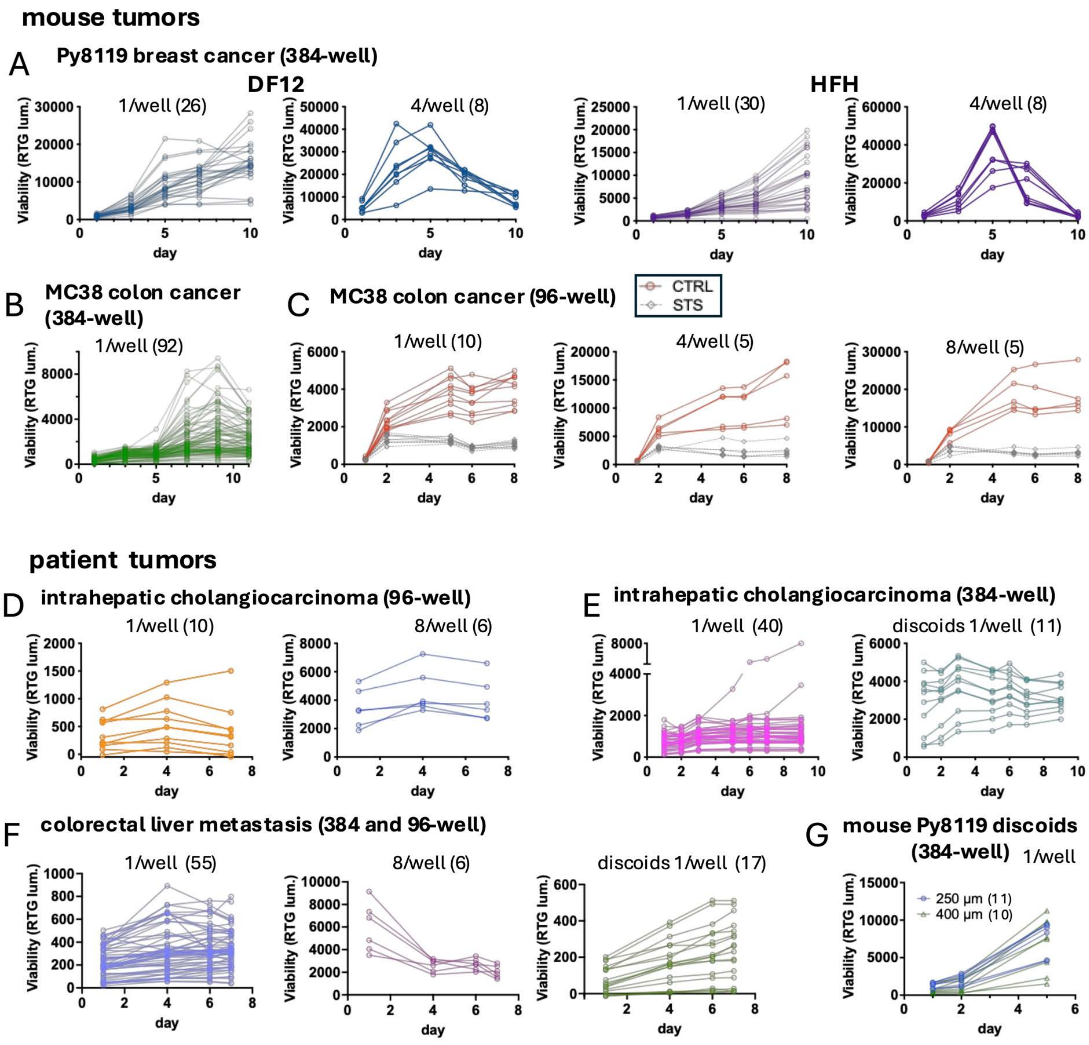


**S1 Fig. Microtumor viability over time.** Cuboids or discoids transferred by a robot into well plates (384 or 96-well as indicated). Viability was measured by RealTime-Glo luminescence on a plate reader, with number of wells in parentheses. Growth medium is HFH unless otherwise indicated. (**A**) Mouse Py8119 breast cancer cuboids grown in DMEM/F12 + 5% heat-inactivated FBS (DF12) or serum-free HFH medium. (**B**) Cuboids from a mouse MC38 colon cancer tumor grown at one per well in a 384-well plate. (**C**) Cuboids from another MC38 tumor grown with different numbers of cuboids/well in a 96-well plate, treated with or without 2 µM staurosporine (STS) starting at day 1. After the day 5 reading, there was a 50% medium change. (**D**) Patient intrahepatic cholangiocarcinoma (ICC) cuboids grown with 1 or 8 cuboids per well of a 96-well plate. (**E**) Individual cuboids and discoids from another patient's ICC grown in a 384-well plate. (**F**) Cuboids and discoids from a patient colorectal liver metastasis. (**G**) Mouse Py8119 discoids cut at either 250 or 400 µm thickness from an 18G needle biopsy, grown in DF12.

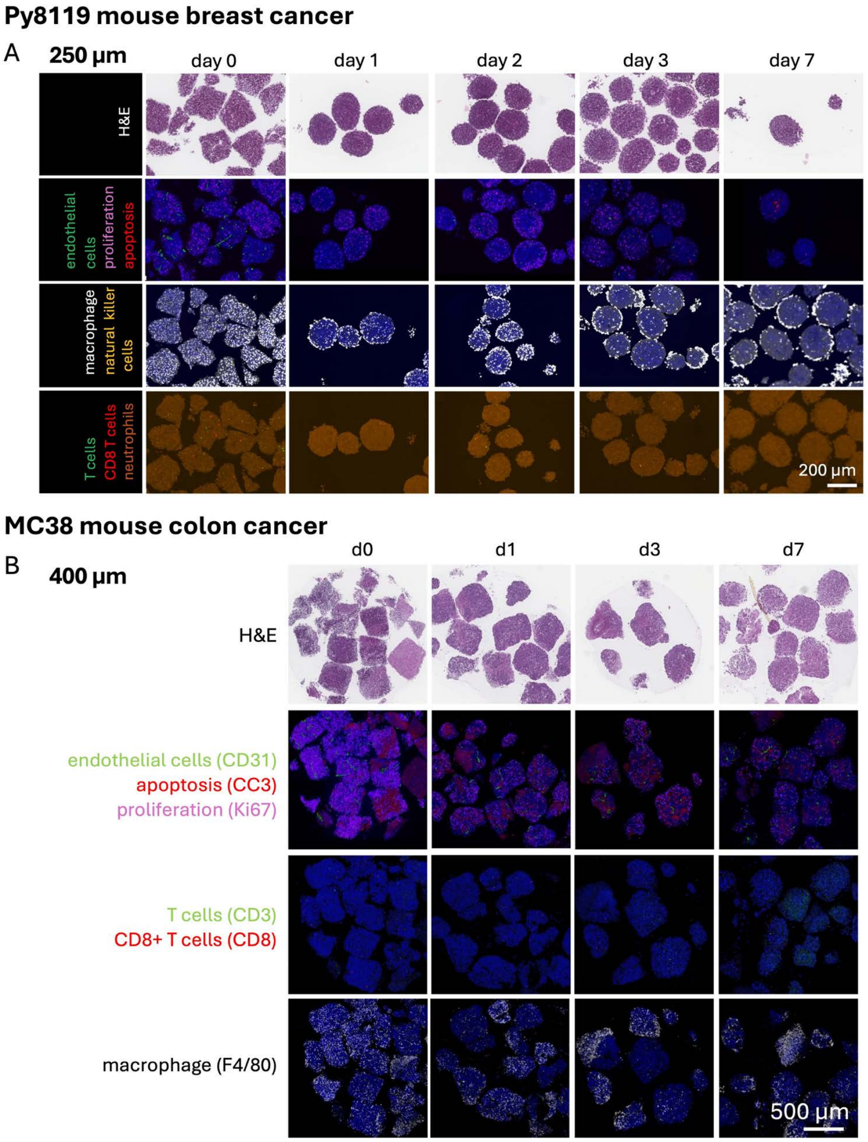


**S2 Fig. Further characterization of the TME of mouse cuboids.** (**A**) Representative brightfield (H&E) and fluorescent multi-immunohistochemistry (multi-IHC) images of 250-µm Py8119 orthotopic mouse breast cancer cuboids from one of two tumors analyzed at different timepoints over 1 week in culture, from the same tumor as in Fig. 2A. (**B**) Histology and multi-IHC images of 400-µm cuboids from a mouse MC38 subcutaneous colon cancer grown in a 6-well plate. Apoptosis (CC3), proliferation (Ki67), endothelial cells (CD31), immune cells (CD45), macrophage (F4/80), NK cells (NKp46), T cells (CD3), cytotoxic T cells (CD8), and neutrophils (Ly6G).

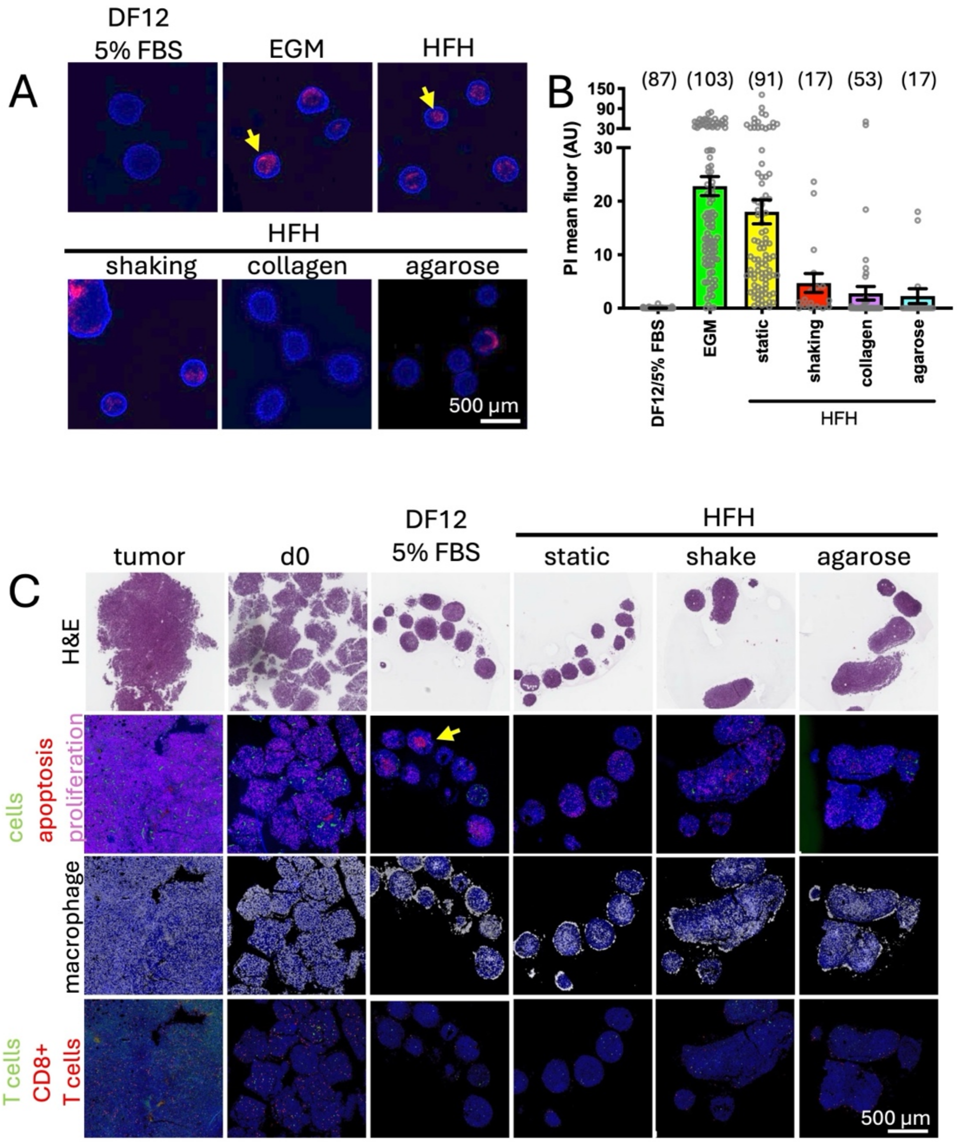


**S3 Fig. Mouse breast cancer cuboids in different culture conditions.** Mouse Py8119 subcutaneous mouse breast cancer cuboids were cultured for 2 days with 50-100 cuboids per well of a 6 well plate media containing serum (DMEM/F12 + heat-inactivated FBS; DF12) or without serum (endothelial growth medium, EGM; or HFH). Some were embedded in 3 mg/ml collagen or in 1% lomelt agarose. (**A**) Cell death at 2 days was measured using the membrane-impermeant red nuclear dye, propidium iodide (PI). Examples of central cell death indicated with yellow arrow. Note that in the shaking conditions, many of the cuboids merged. (**B**) Graph of cell death (PI mean fluorescence). Individual, separated cuboids only. AVE $\pm$ SEM. N in parentheses. (**C**) Histology and multi-IHC at day 0 and day 2 of select conditions as available. Apoptotic cell death (cleaved-caspase 3, CC3), proliferation (Ki67), and multiple cell types. Endothelial cells (CD31), immune cells (CD45), macrophage (F4/80), T cells (CD3), and cytotoxic T cells (CD8). Example of central apoptotic cell death (yellow arrow).

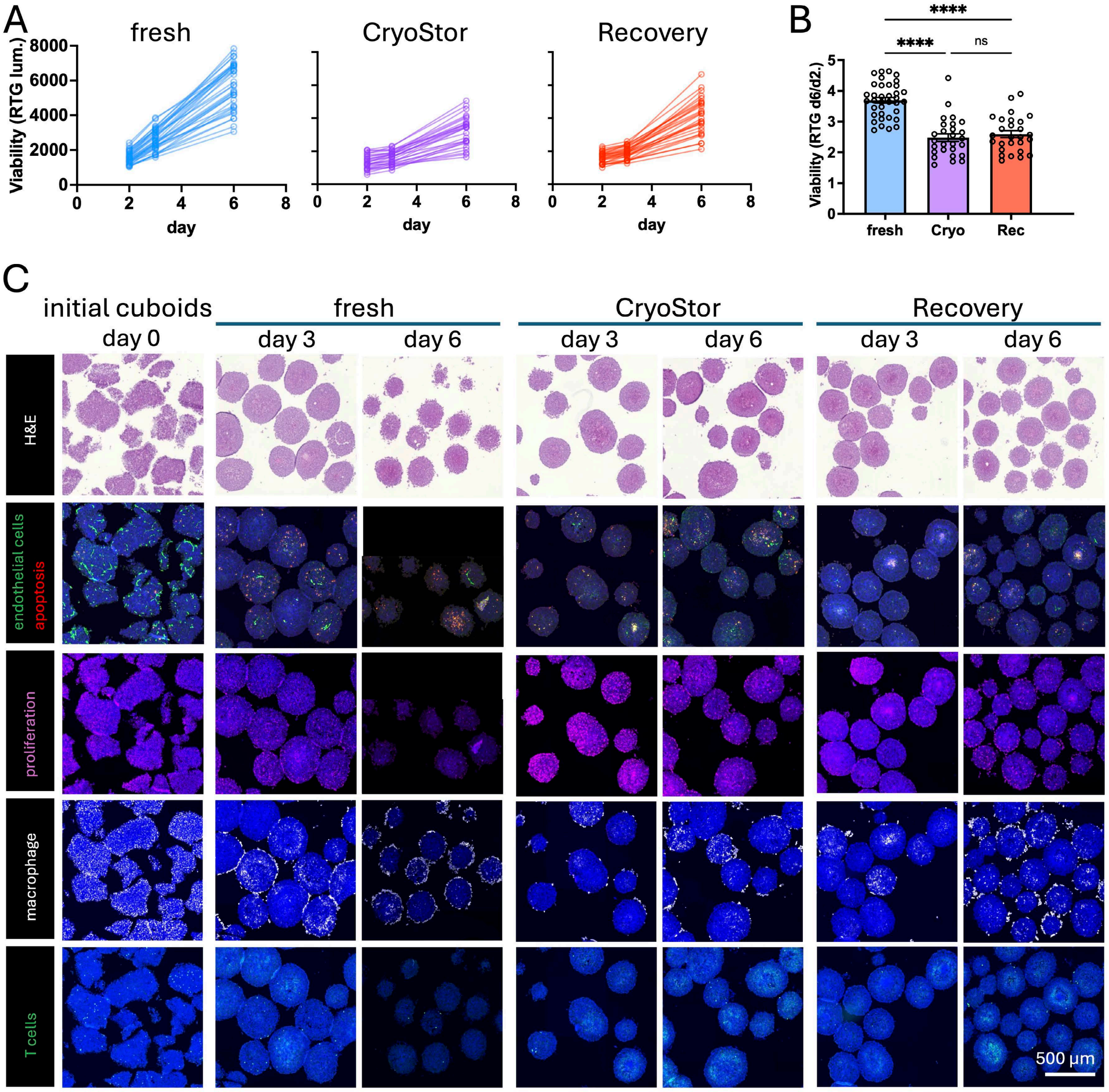


**S4 Fig. Cryopreservation of mouse cuboids.** (**A**) Cuboids from a Py8119 mouse breast cancer, robotically transferred to a 96 well plate, cultured in HFH medium as 4 cuboids/well, either directly (fresh) or after cryopreservation with one of two cryopreservation media (Cryostor or Recovery). Viability was assessed using Realtime-Glo (RTG). n=36 wells for fresh and n= 27 for cryopreserved. (**B**) Change in viability, day 6/day 1. AVE ± SEM. ANOVA with Tukey's post hoc. ****p<0.0001. (**C**) Histology (H&E) and immunostaining of cuboids at different time points for the different treatments. Apoptotic cell death (cleaved-caspase 3, CC3), proliferation (Ki67), endothelial cells (CD31), macrophage (F4/80), and T cells (CD3).

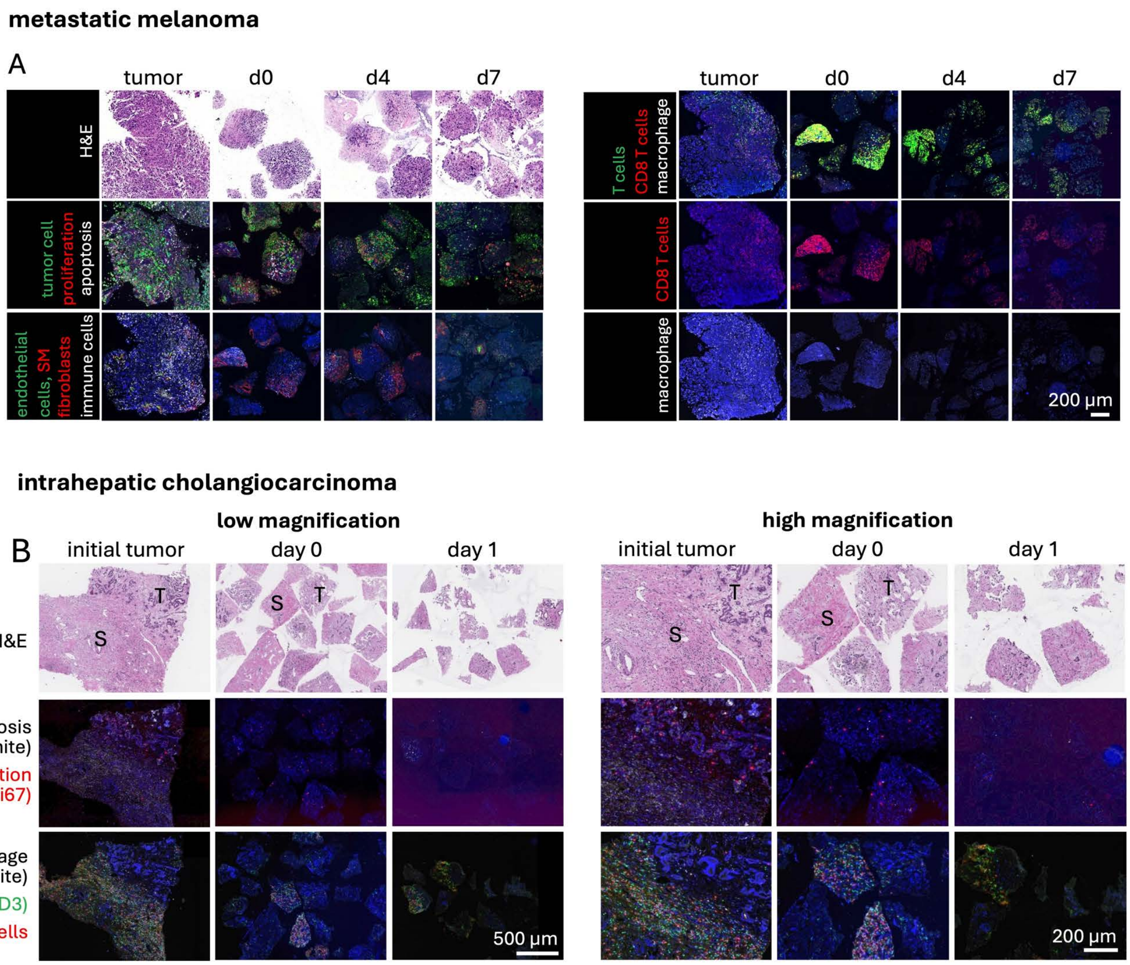


**S5 Fig. Multi-IHC of patient cuboids.** (**A**) Representative images of histology and fluorescent multi-immunohistochemistry (multi-IHC) of cuboids from metastatic melanoma in a lymph node. DAPI nuclear stain in blue. Stain mix 1: Tumor cells (Mel mix), endothelial cells (CD31), fibroblasts and smooth muscle cells (smooth muscle actin), immune cells (CD45). Stain mix 2: macrophage (CD68), T cells (CD3), and cytotoxic T cells (CD8). (**B**) Representative images of histology and multi-IHC of cuboids from an intrahepatic cholangiocarcinoma before and after 1 day in culture, seen at lower and higher magnification. Areas of stroma (S) and of tumor (T) are evident.

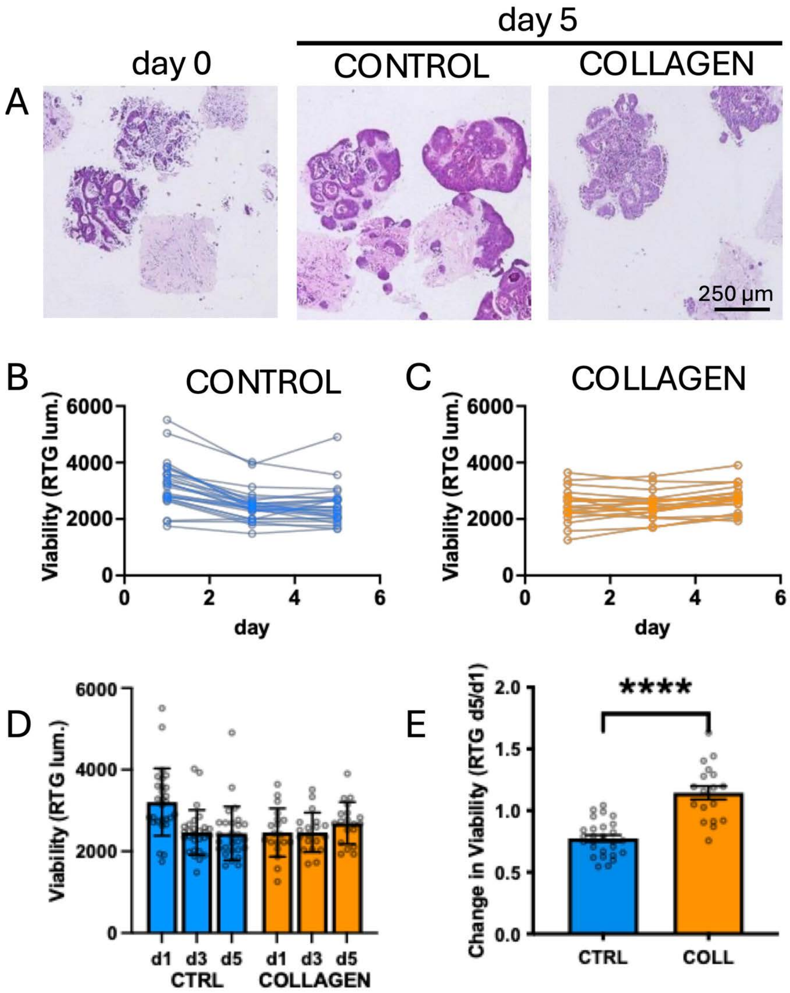


**S6 Fig. Study of patient cuboids cultured in collagen.** (**A**) H&E sections of patient colorectal liver metastasis cuboids were cultured with 3-5 cuboids per well in 96 well plates with collagen (COLL, 50 ul, 2.6 mg/ml, n=18 wells) or without (control, CTRL, n=28). (**B,C**) Viability by RTG over time. (**D**) Summary bar graph of RTG viability over time. AVE ± SD. (**C**) Graph of change in viability from day 1 to day 5 (ratio d5/d1). AVE ± SEM. Student's T-test. ****p<0.0001.

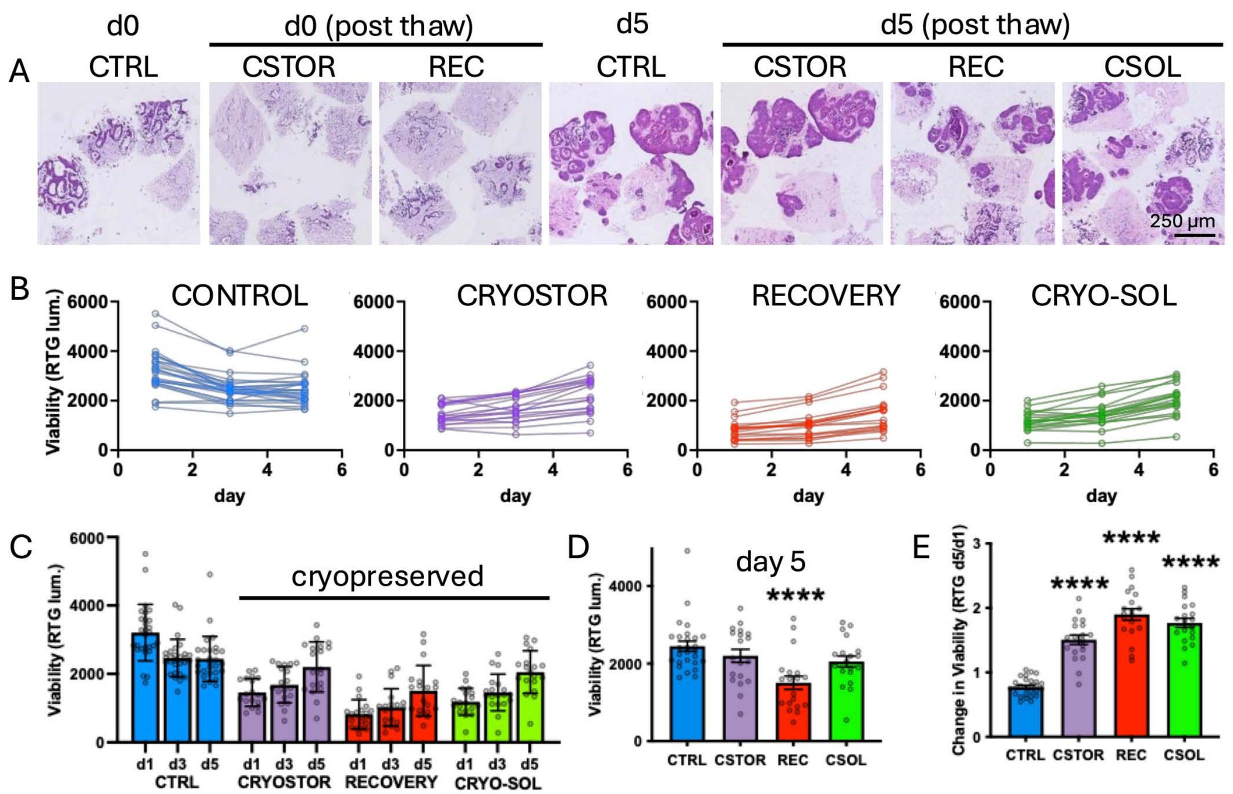


**S7 Fig. Cryopreservation of patient cuboids.** (**A**) H&E histology of patient colorectal liver metastasis cuboids were cultured with 3-5 cuboids per well in 96 well plates either after cryopreservation in different (CryoStor/CSTOR), Recovery/REC, or CRYO-SOL/CSOL; each n=19) or without (control, CTRL, n=28). Day 0 (d0) is the day culture was started, when freshly prepared (CTRL) or after thaw (cryopreserved). (**B**) Viability measured by RealTime-Glo for individual wells over time. (**C**) Summary bar graph of RTG viability with individual wells (points) and AVE ± SD. (**D**) Comparison of RTG viability at day 5. (**E**) Change in viability from day 1 to day 5 (ratio d5/d1). AVE ± SEM. ANOVA versus CTRL with Dunnett's post hoc. ****p<0.0001.

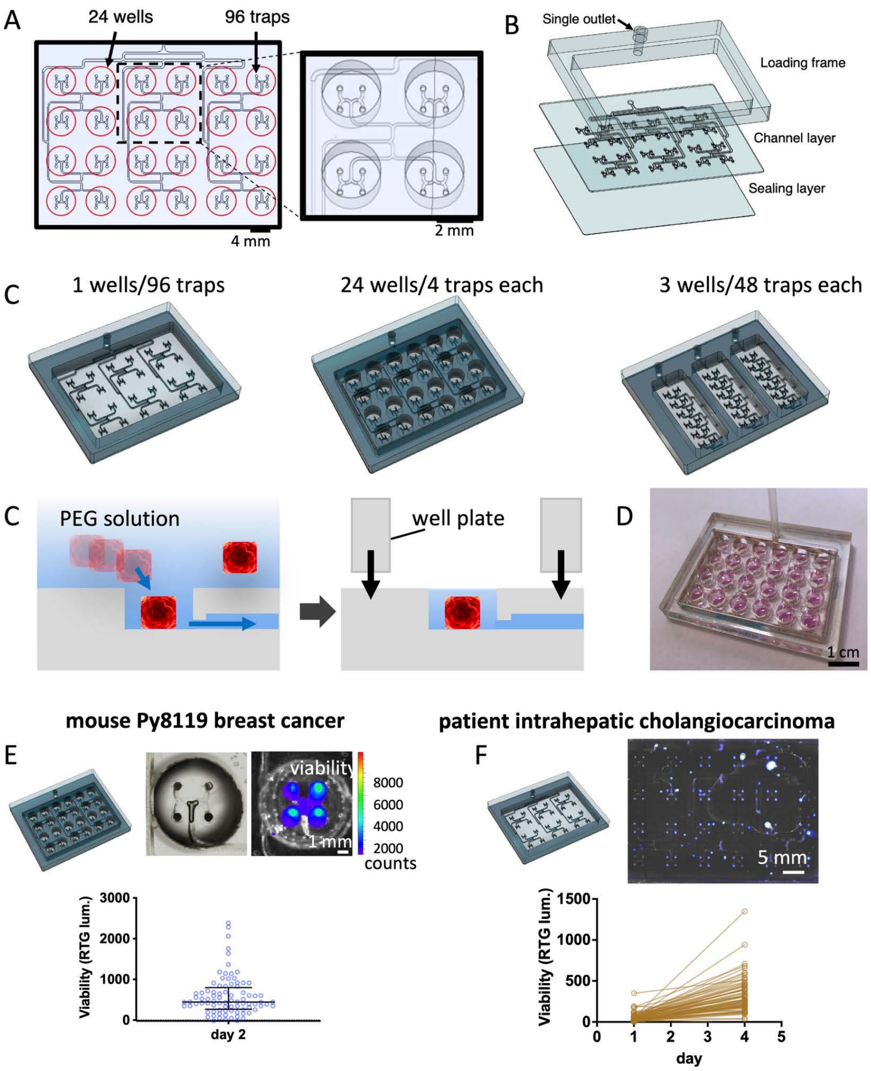


**S8 Fig. Microfluidic device with hydrodynamic traps for 400-µm cuboids.** (**A**) Design of microfluidic device with an array of hydrodynamic traps connected to a single outlet. Red circles indicate location of well from the optional well plate layer that may be added to the device after cuboid loading. (**B**) CAD drawing of the three layers assembled to make the device not including the optional 24 well-plate insert. (**C**) Diagrams of three versions of the microfluidic device. (**D**) Schematic of cuboid loading aided by hydrodynamic trapping (not to scale). Cuboids are either pushed in bulk over the traps or manually approximated to the traps individually for finer control. (**E**) Photograph of an assembled device. (**E,F**) Cuboids grown in microtrap devices with viability measured by RealTime-Glo (RTG) luminescence with a IVIS machine. (**F**) Py8119 orthotopic mouse breast cancer cuboids in a device with 24 wells (4 traps per well), 1-4 cuboids per trap. Schematic of device (upper left), images of one well, brightfield as well as an overlay of RTG luminescence over a photograph (upper right), and graph of viability on day 2 (mean RTG luminescence). n = 77 traps. (**G**) Patient intrahepatic cholangiocarcinoma cuboids grown in a microtrap device with 96 microtraps and only single well (no additional well plate). Manual loading of 1 cuboid per trap. CAD drawing of device (upper left), luminescence image (upper right), and graph of viability (mean luminescence). n = 76 traps.

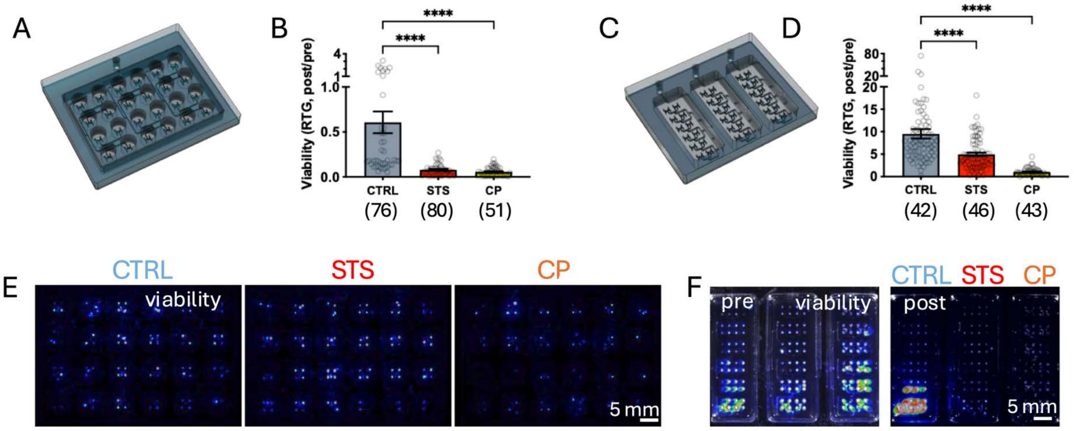


**S9 Fig. Drug responses of 400-µm cuboids from Py8119 orthotopic mouse breast cancers on microdevices.** Cuboids from different tumors were cultured in devices with either 24 wells (96 traps) connected to one outlet (**A,B,E**; 3 devices with 1 drug each) or 3 independent wells with 48 traps each (**C,D,F**; 1 device with 3 drugs). There were 1-4 cuboids per trap. A 3 day drug treatment with cisplatin (100 µM, CP) or staurosporine (2 µM, STS) was started after a baseline reading on day 2. (**A,C**) CAD drawings of devices. (**B,D**) Graph of viability by RealTime-Glo luminescence as post (day 5) versus pre (day 2) drug. AVE ± SEM, individual traps as circles.. Day 2 and 5 measurements taken on different IVIS machines. Kruskal-Wallis test with Dunn's multiple comparisons test versus control (CTRL). ****p<0.0001. (**E,F**) Luminescence images taken by IVIS on day 5 (**E**) or on days 2 (pre) and 5 (post) (**F**).

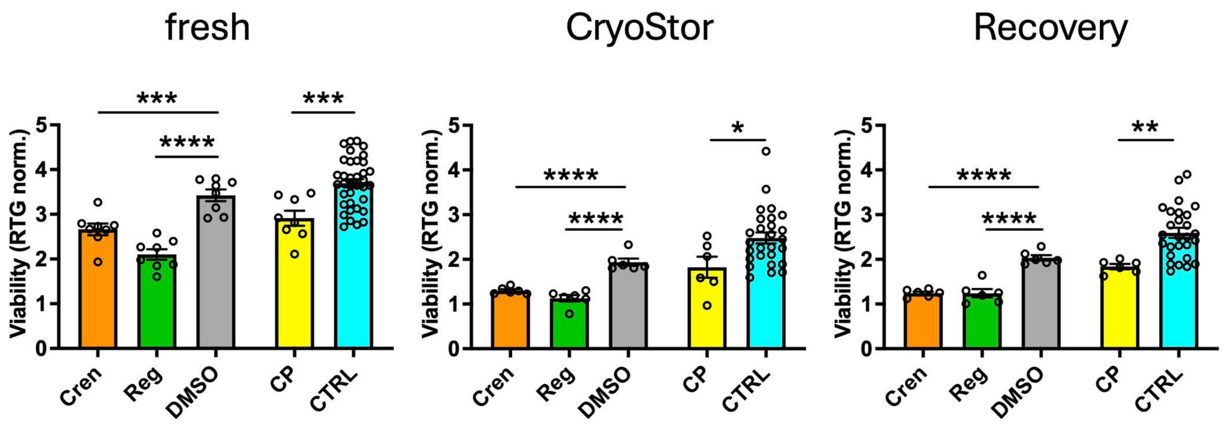


**S10 Fig. Drug testing of cryopreserved mouse breast cancer cuboids.** Cuboids from a Py8119 mouse breast cancer, robotically transferred to a 96 well plate, cultured in HFH medium as 4 cuboids/well, either directly (fresh) or after cryopreservation with one of two cryopreservation media (CryoStor or Recovery). Drug treatment was for 4 days (days 2-6). Drug responses by RealTime-Glo viability (post/pre). Crenolib (Cren 1 µM), Regorafenib (Reg 1 µM) and their vehicle control, DMSO (0.1%). Cisplatin (CP, 100 µM) or control (CTRL). AVE ± SEM. N=6 wells except for controls (CTRL, n=36 for fresh and n= 27 for cryopreserved). One-way ANOVA versus DMSO with Dunnett's post-hoc, or student T-test (CP v CTRL). *p=0.027, **p=0.004, ***p<0.001, ****p<0.0001. Viability, histology, and immunostaining of control cuboids shown in Fig. S4.

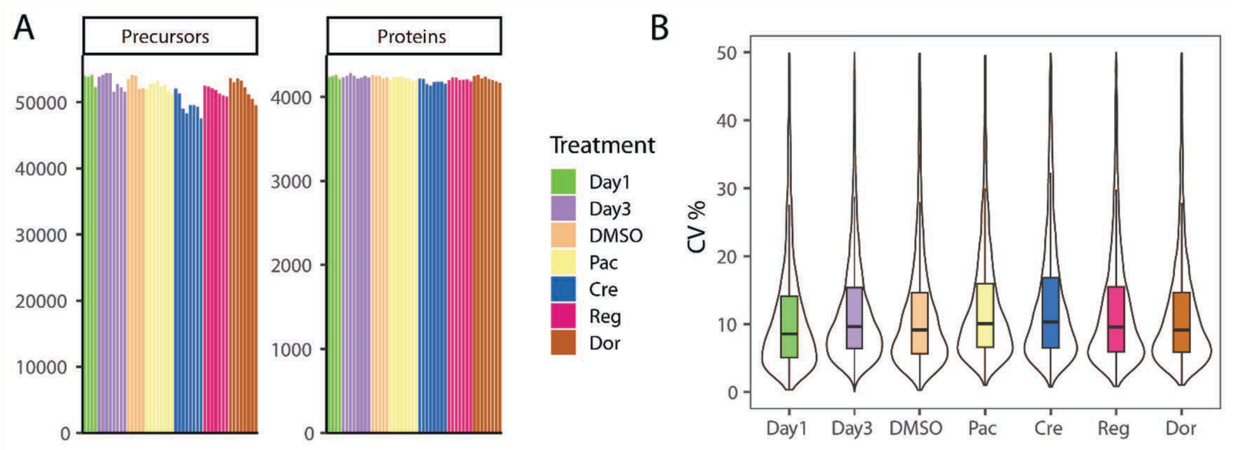


**S11 Fig. Quantification of performance and coverage of cuboid proteome using mass spectrometry proteomics.** (**A**) MSFragger precursor and protein identification statistics for Py8119 breast cancer cuboids across treatment conditions: **Day1**, **Day3**: 0.5 untreated-day 3, **DMSO**: 0.1% DMSO-day 3, **Pac**: 10 µM paclitaxel-day 3, **Cre**: 0.5 µM crenolanib-day 3, **Reg**: µM regorafenib-day 3, **Dor**: 0.5 µM doramapimod-day 3). (**B**) Box Plot and Violin plots: coefficients of variation for normalized protein intensities across treatment replicates.

**S1 Table. Antibody panels used for multi-immunohistochemistry.**

| | Position | Antibody | Clone/Host | Manufacturer/ Catalog Number | Dilution/ concentration | Opal Dye |
|---|---|---|---|---|---|---|
| **mouse immune panel** | 1 | CD45 | IBL-3/16/Rat | Bio-Rad/MCA1388 | 0.181 | **Opal 570** |
| | 2 | CC3 | D3E9/Rabbit | Cell Signaling /9579 | 0.389 | **Opal 690** |
| | 3 | CK19 | TROMA-III/Rat | Developmental Studies Hybridoma Bank (aka DSHB)/TROMA III | 0.736 | **Opal 620** |
| | 4 | CD31 | D8V9E/Rabbit | Cell Signaling/ 776995 | 0.736 | **Opal 520** |
| | 5 | Ki67 | D3B5/Rabbit | Cell Signaling/12202 | 0.736 | **Opal 780** |
| **mouse general panel** | 1 | NKp46 | NCR1/Goat | R&D Systems/AF2225 | 0.389 | **Opal 570** |
| | 2 | CD8a | D4W2Z/Rabbit | Cell Signaling/ 98941 | 2.125 | **Opal 690** |
| | 3 | Ly6g | 1A8 /Rat | Biolegend /127602 | 1.431 | **Opal 620** |
| | 4 | CD3 | EPR 20752 / Rabbit | Abcam/ ab215212 | 1:10,000 | **Opal 520** |
| | 5 | F4/80 | D2S9R/Rabbit | Cell Signaling/70076 | 0.215 | **Opal 780** |
| **human colorectal cancer panel** | 1 | AE1/AE3 | (AE1/AE3)/ Mouse | Dako/M3515 | 0.215 | **Opal 520** |
| | 2 | CC3 | D3E9/Rabbit | Cell Signaling/ 9661 | 0.389 | **Opal 620** |
| | 3 | Ki67 | MIB1/Mouse | Dako/M7240 | 0.111 | **Opal 690** |
| **human melanoma panel** | 1 | Melanoma Cocktail with Sox 10 | Melanoma Cocktail: M2-7C10 + M2-9E3 + T311 + HMB45/Mouse; Sox10: Polyclonal/ Rabbit | Melanoma Cocktail: Novus Biologicals/NBP2-34337; Sox 10: Cell Marque/383A-74 | Melanoma Cocktail: 1:800; Sox 10: 1:50 | **Opal 520** |
| | 2 | CC3 | D3E9/Rabbit | Cell Signaling/ 9661 | 0.389 | **Opal 620** |

| | | | | | | |
|---|---|---|---|---|---|---|
| | 3 | Ki67 | MIB1/Mouse | Dako/M7240 | 0.111 | **Opal 690** |
| **human immune panel** | 1 | CD3 | SP7/Rabbit | Thermo Scientific/RM-9107-S | 0.111 | **Opal 520** |
| | 2 | CD68 | PG-M1/Mouse | Dako/M0876 | 0.597 | **Opal 620** |
| | 3 | CD8 | EP334/Rabbit | BioSB/BSB2848 | 0.389 | **Opal 690** |
| **human cell type panel** | 1 | CD45 | Polyclonal/ Rabbit | Abcam/ab10558 | 0.389 | **Opal 520** |
| | 2 | CD31 | JC70A/Mouse | Dako/M0823 | 0.111 | **Opal 620** |
| | 3 | SMA | 1A4/Mouse | Dako/M0851 | 0.389 | **Opal 690** |

**S2 Table. RPPA raw data for 3 day treatment of Py8119 mouse cuboids.**

| | Phospho-Src Family | Phospho-S6 Ribosomal Protein (Ser240/244) | Phospho-MARCKS | LDHA | Phospho-Myosin Light Chain 2 | Phospho-p44/42 MAPK | Phospho-Bad | Phospho-Stat3 | Phospho-CREB | Phospho-S6 Ribosomal Protein (Ser235/236) | Phospho-Rac1/cdc42 | Phospho-Met |
|---|---|---|---|---|---|---|---|---|---|---|---|---|
| CTL | 91.7166382 | 88.5097447 | 150.4419043 | 106.45 | 97.900887 | 99.08293 | 87.5135554 | 79.12192 | 131.1221 | 96.6035232 | 109.117098 | 112.5582 |
| CTL | 106.977716 | 109.070124 | 113.330754 | 106.16 | 105.056815 | 97.02811 | 94.424563 | 94.48165 | 125.5941 | 76.8250329 | 86.6197506 | 104.1413 |
| DMSO | 131.67404 | 114.984872 | 119.024792 | 109.76 | 131.005666 | 140.6184 | 125.571511 | 151.074 | 145.9553 | 87.285835 | 83.7908166 | 130.4421 |
| DMSO | 68.3259597 | 85.0151283 | 80.975208 | 90.237 | 68.9943335 | 59.38157 | 74.4284893 | 48.92596 | 54.04474 | 112.714165 | 116.209183 | 69.55788 |
| DOX | 0 | 35.8332855 | 0 | 155.1 | 0 | 0 | 32.6958796 | 32.91433 | 0 | 50.2632124 | 74.0773973 | 0 |
| DOX | 0 | 32.1928104 | 0 | 107.51 | 19.7763881 | 0 | 50.3983813 | 110.1415 | 0 | 44.7413701 | 0 | 0 |
| CP10 | 52.7378084 | 64.9519897 | 95.73479613 | 114.81 | 62.9089806 | 38.11904 | 64.8851513 | 6.728785 | 101.1243 | 70.5173325 | 102.92964 | 52.57192 |
| CP10 | 0 | 37.0736121 | 0 | 107.8 | 35.547474 | 0 | 54.4526683 | 124.3827 | 17.22249 | 46.2472208 | 16.9971621 | 0 |
| CP100 | 0 | 0.55866477 | 0 | 156.19 | 0 | 0 | 0 | 27.86253 | 0 | 0 | 8.81951521 | 0 |
| CP100 | 187.54245 | 0.52663332 | 0 | 138.16 | 10.820317 | 0 | 2.38022832 | 4.499059 | 0 | 0.68646861 | 75.9527845 | 0 |
| STS | 0 | 0.54264905 | 0 | 76.358 | 0 | 0 | 0 | 84.74739 | 0 | 37.5214029 | 0 | 0 |
| STS | 0 | 0.54264905 | 0 | 126.76 | 0 | 0 | 0 | 81.97878 | 0 | 0 | 8.58724524 | 0 |

| | Phospho-RSK3 | β-Catenin | Pyruvate Dehydrogenase | Phospho-Pyk2 | Phospho-CaMKII | Phospho-FGF Receptor | Phospho-MDM2 | Phospho-GSK-3β | Phospho-cdc2 | Phospho-Histone H2A.X | PD-L1 |
|---|---|---|---|---|---|---|---|---|---|---|---|
| CTL | 158.303103 | 86.8652546 | 88.00960871 | 158 | 203.414905 | 148.9378 | 122.214229 | 129.3415 | 113.4935 | 106.518432 | 110.736387 |
| CTL | 114.809565 | 72.9770325 | 101.8455998 | 144.23 | 183.024903 | 82.13038 | 105.298734 | 125.1032 | 73.25277 | 96.4934292 | 89.2308321 |
| DMSO | 150.863614 | 105.86769 | 108.7224827 | 131.72 | 91.6216327 | 97.70035 | 104.049898 | 141.87 | 91.98334 | 100 | 114.666577 |
| DMSO | 49.1363865 | 94.1323097 | 91.27751729 | 68.276 | 108.378367 | 102.2996 | 95.9501018 | 58.12996 | 108.0167 | | 85.3334235 |
| DOX | 0 | 27.5096158 | 0 | 0 | 0 | 94.64595 | 80.8896694 | 107.4367 | 130.2979 | 395.469733 | 28.4815687 |
| DOX | 0 | 30.4259326 | 0 | 0 | 0 | 91.37025 | 34.7129406 | 210.6852 | 89.49541 | 417.736835 | 14.2382295 |
| CP10 | 0 | 78.030604 | 78.32993419 | 93.445 | 115.048249 | 101.9203 | 137.562261 | 22.0722 | 158.1908 | 272.010389 | 61.9539556 |
| CP10 | 0 | 52.5591336 | 0 | 0 | 0 | 76.18639 | 84.1780283 | 264.1452 | 133.4944 | 573.648019 | 50.1642826 |
| CP100 | 0 | 0 | 0 | 0 | 0 | 81.98065 | 0 | 100.3145 | 31.68496 | 31.6188558 | 29.7701317 |
| CP100 | 0 | 0 | 0 | 0 | 0 | 94.2824 | 3.98038198 | 14.88337 | 110.4345 | 89.5410571 | 36.2829344 |
| STS | 0 | 0 | 0 | 0 | 0 | 52.91062 | 0 | 122.2584 | 23.95855 | 108.346184 | 5.12641776 |
| STS | 0 | 0 | 0 | 0 | 0 | 64.76735 | 0 | 26.51839 | 69.37454 | 118.765835 | 17.9590606 |

CTL (control), DOX (0.1% DMSO), CP10 (10 µM cisplatin), CP100 (100 µM cisplatin), STS (2 µM staurosporine).

**S1 Movie. Mouse Py8119 mouse breast cancer cuboid day 0.** Whole-mount immunostaining for immune cells (CD45) in red and for endothelial cells (CD31) in green. 3D rendering of confocal images by Imaris.

**S2 Movie. Mouse Py8119 mouse breast cancer cuboid day 4.** Cuboids cultured and processed in microtraps. Whole-mount immunostaining for immune cells (CD45) in red and for endothelial cells (CD31) in green. 3D rendering of confocal images by Imaris.

**S3 Movie. Human intrahepatic cholangiocarinoma cuboid day 0.** Whole-mount immunostaining for immune cells (CD45) in red and apoptosis (CC3) in yellow, with blue DAPI nuclear staining. 3D rendering of confocal images by Imaris.

**S4 Movie. Human intrahepatic cholangiocarinoma cuboid day 4.** Cuboids cultured and processed in microtraps. Whole-mount immunostaining for immune cells (CD45) in red and apoptosis (CC3) in yellow, with blue DAPI nuclear staining. 3D rendering of confocal images by Imaris.
